# Normal and cancer tissues are accurately characterised by intergenic transcription at RNA polymerase 2 binding sites

**DOI:** 10.1101/2023.03.24.534112

**Authors:** Pierre de Langen, Fayrouz Hammal, Elise Guéret, Lionel Spinelli, Benoit Ballester

## Abstract

Intergenic transcription in normal and cancerous tissue is pervasive and incompletely understood. To investigate this activity at a global level, we constructed an atlas of over 180,000 consensus RNA Polymerase II (RNAP2) bound intergenic regions from more than 900 RNAP2 ChIP-seq experiments across normal and cancer samples. Using unsupervised analysis, we identified 51 RNAP2 consensus clusters, many of which map to specific biotypes and identify tissue-specific regulatory signatures. We developed a meta-clustering methodology to integrate our RNAP2 atlas with active transcription across 28,797 RNA-seq samples from TCGA, GTEx and ENCODE, which revealed strong tissue- and disease-specific interconnections between RNAP2 occupancy and transcription. We demonstrate that intergenic transcription at RNAP2 bound regions are novel per-cancer and pan-cancer biomarkers showing genomic and clinically relevant characteristics including the ability to differentiate cancer subtypes and are associated with overall survival. Our results demonstrate the effectiveness of coherent data integration to uncover and characterise intergenic transcriptional activity in both normal and cancer tissues.

## Main

Transcription is a fundamental process in cell biology that converts DNA into biologically active and cell-specific RNA molecules. The majority of transcription is carried out by RNA Polymerase II (RNAP2), which generates mRNAs that are later translated into proteins. However, intergenic active regions have been shown to cover a much larger fraction of the genome than anticipated^1^. Indeed, RNAP2 transcribes a wide variety of intergenic active regions such as many ncRNAs^2^ or enhancer RNAs (eRNAs) that have been found to be major sites of intergenic transcription^3^.

While genes and their protein products have been the main interest in basic and cancer research, an increasing amount of genomic data support the biological and clinical relevance of intergenic transcription. Aberrant expression of non-coding RNAs has been found in cancer^4^ and non-cancer disease^5^, and a large majority of trait or disease-associated variants lie in non-coding regions of the genome^6^. Despite significant progress in describing enhancers transcription^3,7–10^, efforts to fully identify intergenic transcription remains a challenge due to a limited amount of sequencing assays like GRO-seq^11^ or its derivative^12,13^, thus impacting the discovery of a broader intergenic transcription landscape.

In this study, we compiled each available RNAP2 chromatin immunoprecipitation sequencing (ChIP-seq) dataset from GEO^14^ and ENCODE^1^ to construct an atlas of RNAP2 bound intergenic regions in the human genome. Our approach, which targets the RNAP2 rather than the resulting non-coding RNA, aims to minimise the limitation of RNA abundance and stability. This approach enables the exploration of active intergenic regions in a broad range of cell types and tissues, which have not been extensively studied before.

We hypothesise that intergenic RNAP2-bound regions of significance exhibit a biotype-specific signature, reflected in biotype-specific RNA-seq expression across resources such as GTEx^15^, TCGA^16^, and ENCODE^1^. In this study, we describe tissue-specific bindings by creating an atlas of intergenic RNAP2 binding regions. By analysing the expression patterns of 28,797 RNA-sequencing samples, we identify intergenic transcription on RNAP2 binding regions as a powerful indicator for characterising tissue types. We show that using intergenic transcription on RNAP2 binding results in a robust classification of cancer types and subtypes.

Taken together, our study indicates that intergenic transcription at RNA polymerase 2 binding sites is a powerful indicator for characterising normal and cancer tissues at the subtype level. While the functional significance of intergenic regions remains an open question, our findings could significantly enhance our understanding of the regulatory programs and clinical relevance of non-coding transcription in various cancers.

## Results

### An atlas of intergenic RNA Polymerase II occupancy

To create an atlas of intergenic RNA Polymerase 2 (RNAP2) binding in the human genome, we collected all available ChIP-seq targeting RNAP2 on a wide variety of cells and tissue biosamples from public biological data warehouses^1,14^ (**Fig. 1a**). The created atlas aggregates 87% of non-ENCODE datasets and 13% of ENCODE datasets (**Fig. 1a)**. This was accomplished through a standardised manual curation of sample metadata, uniform biosample annotation, and consistent data processing and quality screening, initiated from the raw sequencing files using the ReMap pipeline (Methods). We conservatively retained 906 RNAP2 datasets targeting the POLR2A subunit across 203 cell or tissue types, the majority derived from cancer cell lines (64% of samples) and “normal” cell lines/tissues (36%)(**Fig.1a, Supp Mat, Methods**). In this study, we focused specifically on intergenic RNAP2-bound regions, preventing us from detecting alternative promoters or any transcriptional events occurring within gene bodies (**Methods**). We defined intergenic regions as all regions of the genome, excluding all GENCODE transcripts (as well as known lncRNAs) extended by 1 kb at both TSS and TES, and excluding ENCODE blacklisted region^17^. We identified a total of 23,101,589 RNAP2 binding events across all 906 datasets, of which 2,525,886 (11.1%) are localised within intergenic regions (averaging 2,787 intergenic binding events per dataset, **Supp. Fig. 2a**). A large fraction of RNAP2 intergenic binding events (91.7%) are extensively shared across experiments, indicating similar occupancy patterns **(Fig. 1b)**. We developed an aggregative approach to identify across experiments what we refer to a “Consensus Peaks” (**Fig. 1a, Methods**). By applying this approach, we created an atlas of 181,547 intergenic RNAP2 consensus, describing distinct genomic elements bound by RNAP2 across multiple biosamples. Our atlas of intergenic RNAP2-occupied regions, which is available on Zenodo^18^, is based on consensus peaks derived from 13 datasets on average, with each consensus having an average width of 470 bps (**Fig. 1c** and **Supp. Fig. 2c**). Each peak in the ChIP-seq data contributing to a representative RNAP2 consensus can be traced back to its corresponding biosample (**Supp. Fig. 1b**). We evaluated our created atlas against reference databases of regulatory and non-coding genomic elements **(Fig. 1d,e**). Our analysis revealed that intergenic RNAP2 consensus are preferentially located over regulatory elements and potentially transcribed enhancer regions.

**Fig 1:**
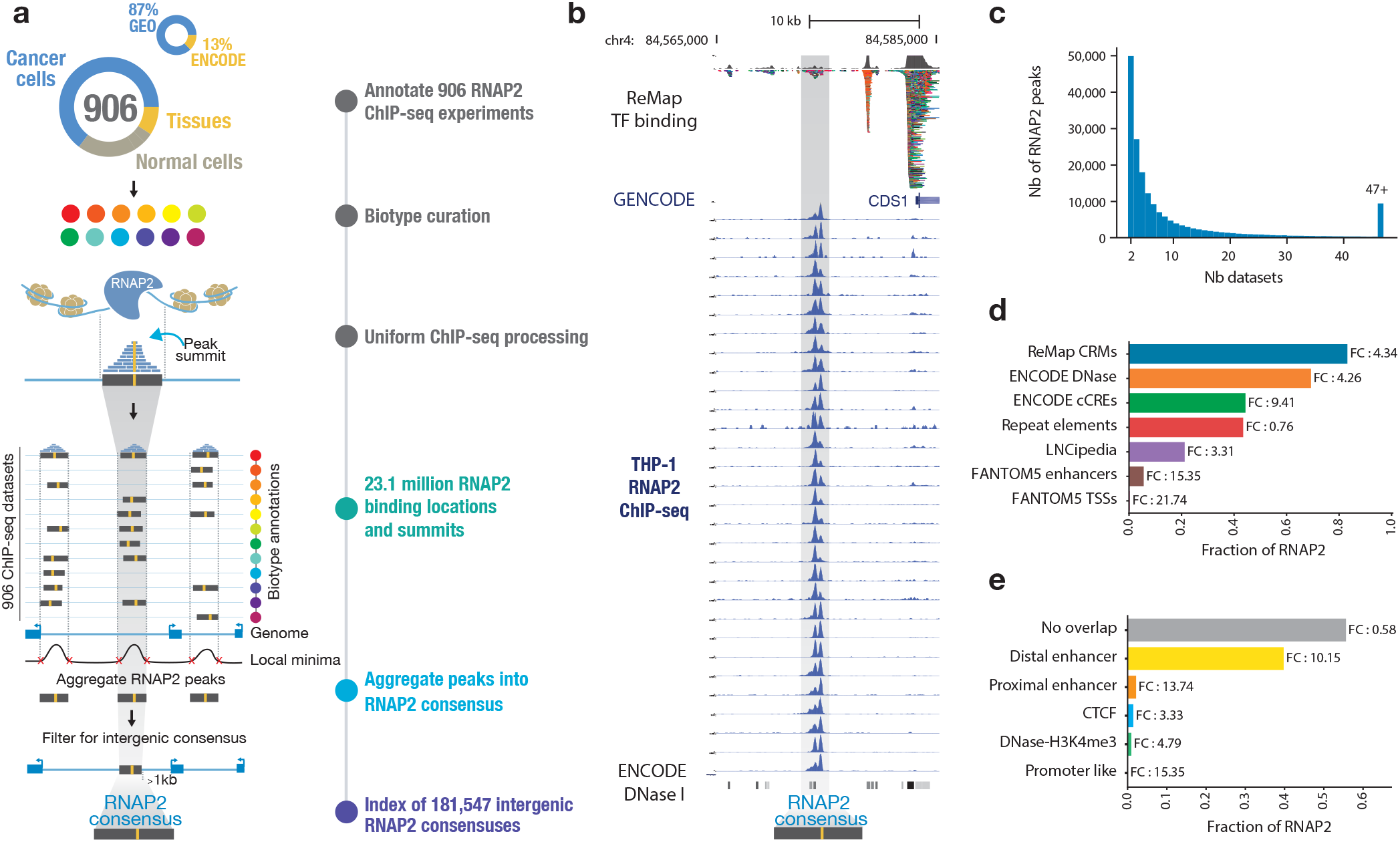
An atlas of intergenic RNA Polymerase II occupancy. **a**, Outline of the RNAP2 atlas pipeline; 23.1 million RNAP2 bound regions aggregated across 906 individual datasets jointly identify 181,547 intergenic RNAP2 consensus. **b**, Example locus on chromosome 4, showing RNAP2 ChIP-seq density across THP-1 cell lines (leukaemia) with location of a RNAP2 consensus. **c**, Distribution of the number of datasets across which RNAP2 peaks are shared. **d**, Comparison of RNAP2 consensus location with resources of regulatory and non-coding elements. FC denotes fold change versus random; all results are statistically significant. **e**, Distribution of ENCODE cCREs within the RNAP2 atlas.

### A normalised vocabulary captures biotype-specific intergenic RNP2 binding

The RNAP2 atlas covers a very large fraction of the human biological spectrum, with over 203 distinct tissues and cell lines (**Fig. 2a, Supp. Table 1**). To facilitate biological interpretation, we grouped biosample annotations based on their tissue of origin or similarity. We then grouped similar tissues into 16 distinct biotypes to obtain a concise yet meaningful high-level annotation of our samples (**Supp. Table 1, Methods**). To simplify genomic interoperability across large resources, the compendium of tissues and cell lines were harmonised using GTEx, TCGA, ENCODE biosamples nomenclature as well as cell ontologies^19,20^. This results in RNAP2 consensus exhibiting a biological context ranging from biotypes-specific to ubiquitous signatures (**Fig. 2b**). As intergenic RNAP2 binding appears extensively shared across biosamples (**Fig. 1b,c**), we aimed to visualise RNAP2 occupancy patterns across biosamples and consensuses by employing a hierarchical clustering approach (**Fig. 2b, Methods**). Patterns of RNAP2 binding were structured into mostly biotype-specific and a few ubiquitous occupancy clusters. We observed seemingly sparse distribution in the intergenic RNAP2 atlas, but upon further investigation, we identified diverse and intricate binding patterns. To analyse these patterns, we utilised an unsupervised dimensionality reduction technique (UMAP^21^) on the 906 biosamples (**Fig. 2c**) and >180,000 RNAP2 consensus (**Fig. 2d**). The UMAP visualisation across 906 ChIP-seq datasets reveals organised intergenic occupancy patterns across similar biotypes (**Fig. 2c**). Based on their intergenic occupancy patterns, datasets having similar biotypes of origin are clustered together, while the centre of the plot contains datasets with ubiquitous biotype signatures. For example, ChIP-seq datasets for digestive biosamples (Brown dots, n=126 samples) were predominantly clustered together, suggesting that intergenic RNAP2 occupancy is representative of the sample biology, but also that the biosample curation is coherent. Next, we visualised the 181,547 intergenic RNAP2 consensus according to their binding patterns and biotype labels (**Fig. 2d, Methods**). To facilitate the biological interpretation of a consensus, each consensus is labelled with their most frequent biotype, or labelled in grey when ubiquitous. By visualising the intergenic RNAP2 atlas, we were able to identify distinct occupancy patterns that are specific to certain biotypes. This framework was also applied on 890 H3K27ac datasets, successfully demonstrating its ability to identify biotype-specific clusters of histone modifications (**Supp. Fig 3**). The RNAP2 atlas, generated by leveraging 906 ChIP-seq datasets, provides a valuable biotype-specific summary of intergenic RNA Polymerase 2 binding. Its potential to uncover intergenic transcriptional activities makes this atlas an innovative tool.

**Fig 2:**
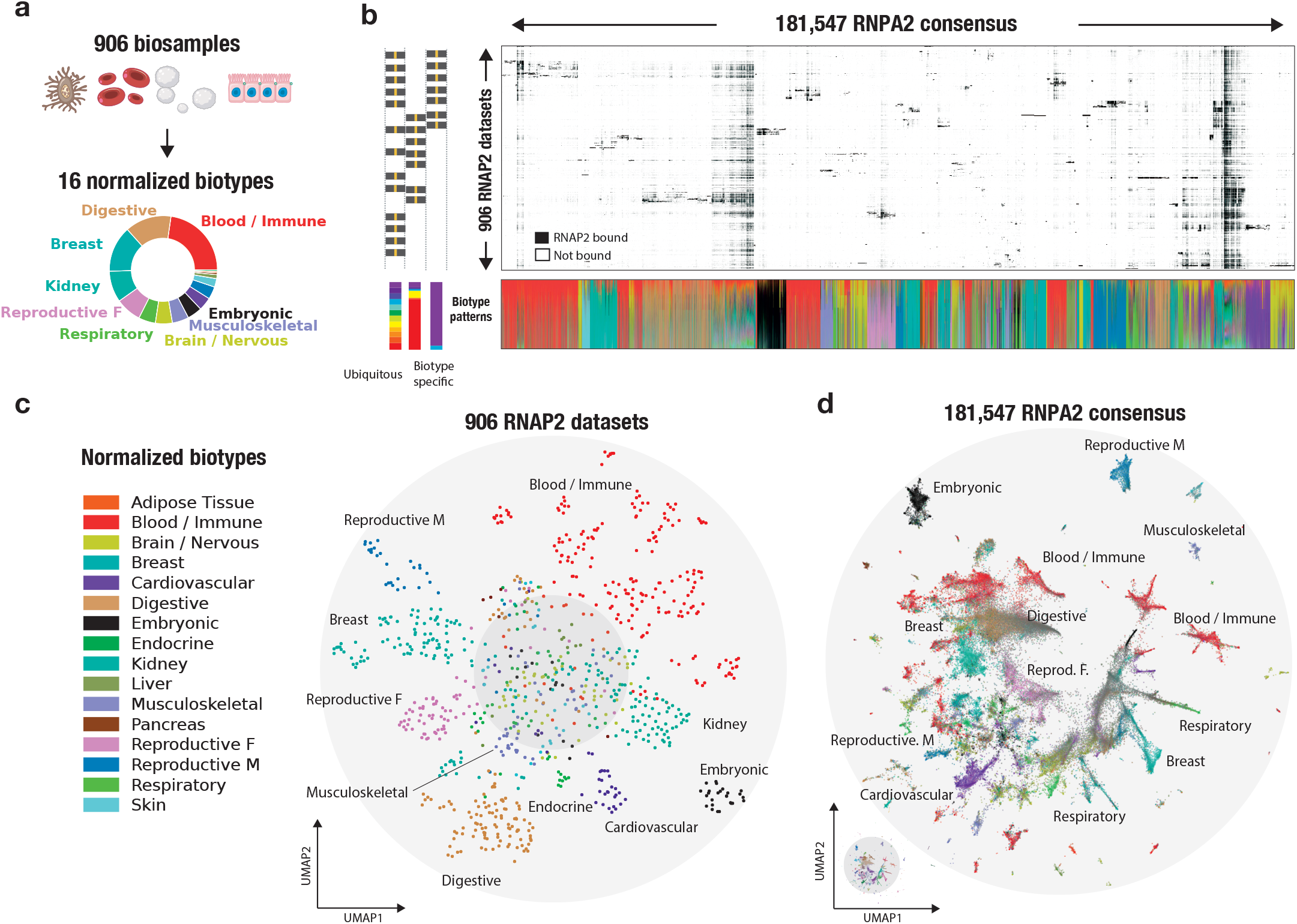
A normalised vocabulary captures biotype-specific intergenic RNP2 binding. **a**, Distinct tissues and cell lines across 906 biosamples normalised into 16 biotypes. **b**, Intergenic RNAP2 occupancy in 181,547 consensus regions across 906 biosamples displayed in a visually compressed matrix. The colour code used for each RNAP2 consensus region corresponds to the biosample tissue of origin, with examples representing either biotype-specific or ubiquitous signatures. This colour scheme is consistently applied across all RNAP2 consensus regions. Lower panel indicates the normalised contribution of a biotype, in terms of peaks, to each RNAP2 consensus (Methods). **c**, Two-dimensional Uniform Manifold Approximation and Projection (UMAP) projection of all 906 RNAP2 ChIP-seq datasets across intergenic RNAP2 space, coloured by normalised biotype. **d**, UMAP projection of all intergenic RNAP2 consensus according to their binding patterns, coloured by dominant biotype (Methods).

### Revealing tissue-specific regulatory signatures

We next sought to retrieve and annotate each consensus group to capture their biological identity. Using an unsupervised graph clustering approach we identified 51 RNAP2 consensus clusters (**Fig. 3a**), each harbouring its own tissue-specificity (**Fig. 3b**). To validate their biological signatures, the clusters were compared against the biological classification of the human index of DNase I hypersensitive sites^22^ (DHSs)(**Fig. 3c, Supp. Fig 5a**). The defined RNAP2 clusters display a coherent enrichment with the DHSs regulatory vocabulary, for example the ‘Brain/Nervous’ RNAP2 cluster #31 (light green) is enriched in Neural DHSs. To capture the genomic signatures of these groups, we examined the epigenetic state per RNAP2 cluster, and more specifically, its chromatin state specificity. As an example, we took the RNAP2 cluster #4 having a clear “Embryonic” signature and sampled the Roadmap ChromHMM epigenetic states of Embryonic Stem Cells (**Fig. 3d**). We observe a strong enrichment of “active” epigenetic states, notably Enhancers, TSSs and transcribed regions within the RNAP2 embryonic cluster compared to the other RNAP2 (**Supp. Fig. 4b, Methods**). On the other hand, a depletion in “inactive” epigenetic states, such as Quiescent or Polycomb-repressed states is observed (**Fig. 3d**), suggesting that RNAP2 occupies intergenic space at key regulatory elements as previously shown^23^ (**Fig. 1d,e**). We explored the tissue specificity of RNAP2 clusters by analysing enhancer-like histone marks and open chromatin profiles (H3K27ac, ATAC-seq, **Supp. Table 2**). The results showed that RNAP2 cluster-tissue pairs with matching tissues (e.g., Heart-Cardiovascular) had the strongest activity, while non-matching pairs (e.g., Lymphoid-Liver) had a weaker signal (**Supp. Fig. 4a**).

**Fig 3:**
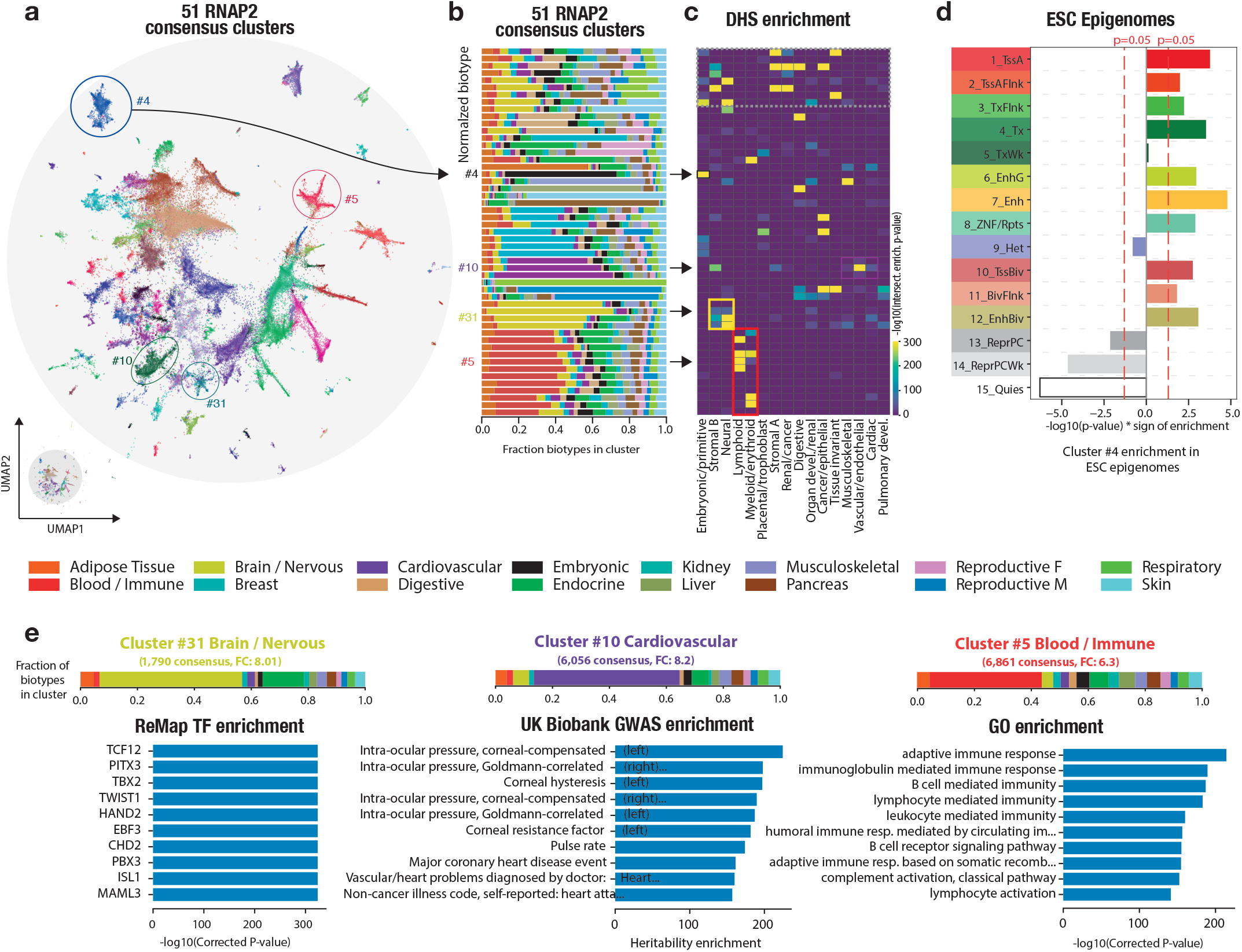
Revealing tissue-specific regulatory signatures. **a**, Unsupervised graph clustering identifies 51 RNAP2 consensus clusters. Four clusters (4, 5, 10, 31) are highlighted across panels a, b, c to illustrate the analysis. **b**, The fraction of biotypes within each cluster is shown, indicating tissue-specific or ubiquitous signatures. **c**, Enrichment of DNase I hypersensitive sites (DHS) biological classification in each cluster is depicted. Arrows and coloured rectangles highlight correspondence between clusters and DHS categories. **d**, Enrichments of ChromHMM epigenetic states of “Embryonic Stem Cell”, sampled at RNAP2 genomic location of Cluster #4, against non-cluster #4 RNAP2 consensus. Active states; active TSS transcription states (TssA, TssAFlnk), transcribed promoter and enhancer signatures (TxFlnk), actively-transcribed states (Tx, TxWk), enhancer states (Enh, EnhG), zinc finger protein genes state (ZNF/Rpts). Inactive states ; heterochromatin (Het), bivalent regulatory states (TssBiv, BivFlnk, EnhBiv), repressed Polycomb states (ReprPC, ReprPCWk), quiescent state (Quies). **e**, Top 10 Transcription factor enrichments from the ReMap database in the cluster #31; Top 10 UK Biobank GWAS trait heritability enrichment in the cluster #10; Top 10 GO enrichment of nearby genes in the cluster #5. All results shown are statistically significant. Each cluster’s biotype distribution is shown as a stacked bar plot.

To further confirm the biological identity of each cluster, we investigated enrichment of SNP-based trait heritability from UK Biobank GWAS^24^, transcription factors binding regions (TFBRs) from ReMap^25^ and Gene Ontology (GO) terms (**Fig. 3e**). The “Cardiovascular” cluster #10 is enriched in multiple heart related traits (Intra-corneal pressure, Pulse rate, Coronary heart disease). The “Brain/Nervous” cluster #31 showed TFBRs enrichment of TFs known to be involved in Neural development, or diseases (TCF12, PITX3, TWIST1). Consistent with their assigned biotypes, the Blood/Immune RNAP2 clusters #5 is located near genes linked to immune response GO terms. Our study accurately distinguishes intergenic RNAP2 occupancy based on its biotype-specificity, revealing tissue-specific regulatory signatures across multiple independent genomic resources, ranging from open chromatin occupancy maps to transcription factor binding.

### Systematic transcription captured in intergenic RNAP2 atlas

We developed the RNAP2 atlas as an innovative tool for indirectly identifying intergenic regulatory regions that are active or poised for transcription. To quantify intergenic transcription and gain a better understanding of transcriptional patterns, we utilised the RNAP2 atlas to probe transcriptional signals in three of the largest expression resources available, which include samples from normal, cancer and cell lines: the Genotype-Tissue Expression (GTEx), the Cancer Genome Atlas (TCGA), and the ENCODE consortium. By combining these resources, we conducted a comprehensive analysis of intergenic expression across the RNAP2 atlas, leveraging data from 28,767 RNA-seq samples. To quantify intergenic transcription, we first standardised each RNAP2 consensus sequence to a 1kb RNAP2-bound region. We then counted the number of reads that overlapped with these RNAP2-bound regions and generated a count table similar to conventional single-cell RNA sequencing (scRNA-seq) count tables (**Fig. 4a**). We found that the intergenic RNAP2 atlas captured approximately 60% of intergenic reads (**Supp. Fig. 8**). Furthermore, these RNAP2-bound intergenic regions captured significantly more reads than the rest of the intergenic genome. On average, RNAP2-bound regions had 7.13 times more transcriptional signals compared to the rest of the intergenic genome (**Fig. 4b, Supp. Fig. 8a**,**b**). Altogether, the RNAP2 atlas is strongly enriched in transcriptional activity, and thus could serve as a powerful tool for investigating intergenic transcription in both normal and cancer tissues.

**Fig 4:**
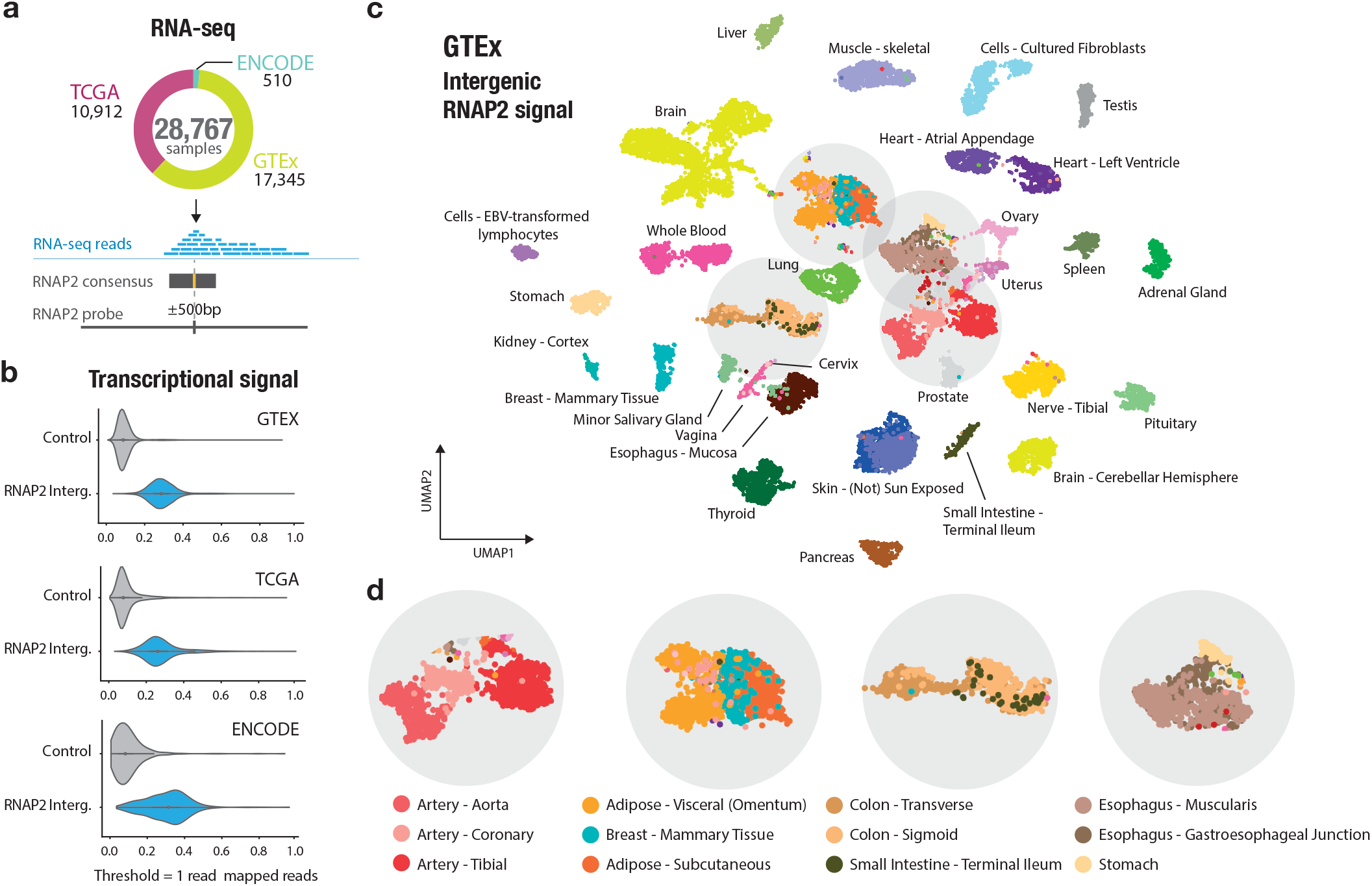
Intergenic transcription on RNAP2 atlas is a powerful indicator for characterising tissues. **a**, Number of RNA-seq samples from three expression resources (GTEx, TCGA, ENCODE) and schema depicting the standardisation of RNAP2 consensus to 1kb “RNAP2 bound regions” to obtain read counts. **b**, Violin plots comparing transcriptional signals at intergenic RNAP2 bound regions versus non-RNAP2 random intergenic regions across the three expression resources. **c**, Two-dimensional UMAP projection of 17,345 GTEx RNA-seq signals across the intergenic RNAP2 atlas, with colours representing 54 tissue types, including 11 distinct brain regions (yellow) and two cell lines (light blue). **d**, Zoomed view of tissue-specific expression patterns observed in similar tissues, such as different types of Artery (e.g. Aorta, Coronary, Tibial).

### Intergenic transcription on RNAP2 atlas is a powerful indicator for characterising tissues

To determine whether intergenic transcription at RNAP2-bound regions could characterise tissue specificity, we analysed expression data from 54 non-diseased tissues across 17,345 samples from the GTEx project. Intergenic transcription has been used as a marker of enhancer activity as described in the FANTOM project^7^, and across the development of experimental assays enriched for capped and nascent RNAs^3,10,12,13^. Here, we have developed a pipeline based on scRNA-seq methods, which commonly deals with weak signals and large sample sizes. By using only signals on RNAP2 bound regions, we were able to extract valuable biological information from read count tables (**Supp. Fig. 6, Methods**). We used UMAP to visualise similarity between the expression levels sampled at RNAP2 bound regions of each GTEx biosample, which revealed a clear distinction between tissues as biosamples from the same sampling site are clustered together (**Fig. 4c**). These tissue-specific expression patterns are observed between similar tissues such as Artery (Aorta, Coronary, Tibial), or between tissues with similar histological features such as Adipose tissues (Visceral, Subcutaneous, Mammary tissue), or at different body locations with samples originating from the digestive tract (Colon, Small intestine) (**Fig. 4d**). To test whether intergenic transcription could discriminate the 54 GTEx tissues accurately, we used a k-nearest neighbours algorithm (KNN classifier) for RNAP2 bound regions and gene centric RNA-seq counts. We found that RNAP2 bound regions could predict tissue types with a high level of accuracy, demonstrating only a slight decrease in accuracy compared to gene-centric RNA-seq counts processed using the same methods (87.1% against 90.0% balanced accuracy across 54 tissues, **Supp. Fig. 9c**). Next, we obtained over-expressed intergenic RNAP2 bound regions in the GTEx tissues, with an average of 4,236 regions per tissue (**Supp. Fig. 10c, Methods**). Our analysis showed a significant association between RNAP2 bound regions with tissue-specific over-expression and tissue-specific GTEx eQTLs (**Supp. Fig. 10a**,**b, Methods**), providing evidence that the RNAP2 bound regions can be used to detect transcribed intergenic enhancers. Interestingly, we detected transcriptional signals at RNAP2 bound regions located downstream genes (>1kb) indicating that there could be transient RNA downstream of the polyadenylation site (**Supp. Fig. 11a**), consistent with previous studies^26^. To investigate the effect of these signals, we performed additional analyses that excluded RNAP2 bound regions located up to 9kb downstream of genes. Our findings demonstrate that RNAP2 consensus located within the 1-9kb tail of genes do not drive the classification of GTEx tissues (**Supp. Fig. 11b**,**c**). Furthermore, we show that the RNAP2 atlas is applicable to smaller RNA-seq samples. We compared expression in three samples of two types of heart tissues from GTEx biosamples and identified 195 RNAP2 bound regions located near genes related to heart functions, despite limited statistical power (**Supp. Fig. 12, Methods**). Here, we provide evidence that intergenic transcription detected at RNAP2-bound regions is a strong indicator of tissue specificity and can be used to accurately predict tissue types. Our results may have implications for understanding tissue-specific gene regulation.

### Meta-analysis reveals tissue-and disease-specific connections between RNAP2 occupancy and transcription

We examined the relationship between biotype-specific RNAP2 occupancy and biotype-specific transcription by comparing the observed intergenic signal across all expression datasets, combining 28,787 RNA-seq samples, despite the use of different sequencing samples and protocols. We first conducted an analysis to determine the relationship between biotype-specific RNAP2 occupancy in ChIP-seq and transcription in ENCODE RNA-seq biotypes by comparing biotypes pairwise **(Fig. 5a**). This revealed a strong biotype-specific transcriptional enrichment in the ENCODE dataset at RNAP2 probes with a ChIP-seq occupancy specific to the same biotype, even through different samples and protocols. On the other hand, non-matching biotype pairs do not show transcriptional signal enrichments. This highlights a strong association between RNAP2 occupancy and effective transcription, as well as the effectiveness of our biosample annotation for varied data sources comparisons. Furthermore, we performed a meta-analysis integrating every dataset and biotype, to obtain a connected, global view of the intergenic transcription across almost 30,000 biosamples of varied data sources. Briefly, we extracted intergenic RNAP2 markers (only up-regulated RNAP2 bound regions, **Methods**) for every possible biotype-dataset pair (i.e. RNAP2-Liver, GTEx-Heart, ENCODE-Liver…), and quantified pairwise similarity between every biotype-dataset marker lists, assuming that a marker list is characteristic of a biotype. Then, we applied Hierarchical Clustering to obtain a meta-clustering revealing similarities between tissues across all resources (**Fig. 5b, Methods**). This meta-analysis highlights that the connection between intergenic RNAP2 occupancy and intergenic transcription is biotype-specific, consistently observed across biotypes, and independent of dataset origins or protocols used. Our approach groups biotypes and similar biotypes together, independently of the data source (**Fig. 5b**). For example, “Adipose tissue” and “Breast” tissues cluster together across resources as breast tissue contains adipose cells (**Fig. 5c**). Globally, identical biotypes also have much more similar markers across data sources than non-identical biotypes (**Fig. 5d**). For each biotype, we extracted robust markers supported by at least half of the data sources. We observe that these markers showed strong heritability enrichment in biotype-related traits, confirming their biological relevance (**Supp. Fig. 13**). For example, “Reproductive Female’’ markers were strongly associated with heritability for the “Birth weight of first child” trait, while “High cholesterol” heritability was enriched within “Liver” markers. In summary, our meta-analysis revealed a tissue-specific, biologically meaningful correlation between intergenic transcription and RNAP2 occupancy, as well as a strong coherence across varied data sources and protocols.

**Fig 5:**
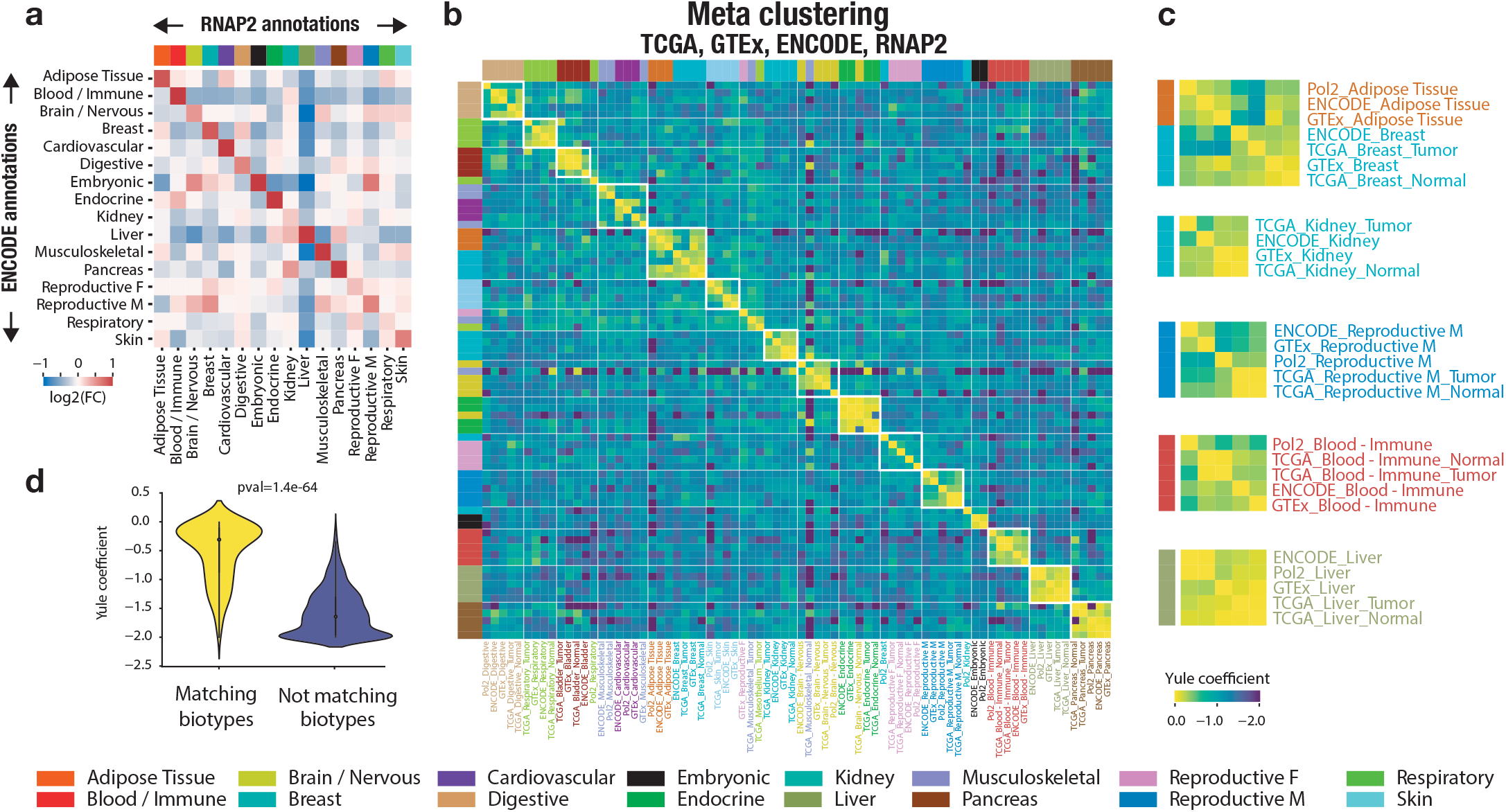
Meta-analysis reveals tissue-and disease-specific connections between RNAP2 occupancy and transcription. **a**, Association between RNAP2 occupancy biotype and transcription biotype from ENCODE. Heatmap depicts Log2 of ENCODE RNA-seq dataset expression Fold Change, in each biotype (rows), between RNAP2 bound regions with biotype-specific RNAP2 ChIP-seq occupancy (columns) against non-specific RNAP2 bound regions. **b**, Heatmap showing the association between biotype-specific intergenic RNAP2 occupancy and biotype-specific RNAP2 over-expression across four resources. Hierarchically clustered heatmap reveals the correct grouping by tissue of origin rather than data source, with each possible biotype-dataset pair represented. Yule distance between a pair of dataset-biotype lists of over-expressed RNAP2 markers is indicated. **c**, Zoomed view revealing meta-clusters of tissue-specific correlation between intergenic RNAP2 regions and their transcription in different ressources. **d**, Distributions of tissue-matching (i.e. “Pol2-Liver” vs “TCGA-Liver”) and non-matching (i.e. “Pol2-Liver” vs “GTEx-Heart”) Yule distance between two intergenic RNAP2 marker sets (p-val=1.4e-64).

### Cancer types and subtypes classification by intergenic transcription at RNAP2 binding sites

We have shown that intergenic transcription can reliably discriminate between various tissues and biological conditions. Building on this insight, we explored potential direct applications of our RNAP2 atlas and approaches to human cancers. We analysed expression data from 32 cancer types across 10,912 RNA-seq samples from the TCGA cohort to identify clinically relevant intergenic transcription and potential therapeutic targets (**Fig. 6a**). We visualised the biosample expression similarity using UMAP, which first revealed a distinction between cancer types, then, by tissue state (normal tissue or tumoral tissue), suggesting that some RNAP2 bound regions are differentially expressed in these contexts (**Fig. 6b**). For example, two brain cancers such as the Lower Grade Glioma (LGG) and Glioblastoma multiforme (GBM) cluster closely together, while Kidney tumour samples (KIRC, KICH, KIRP) have distinct profiles, even if normal kidney samples appear similar. Interestingly, Breast cancer samples (BRCA) cluster into two distinct groups on the basis of intergenic RNAP2 bound regions expression. These two groups correspond to distinct breast cancer subtypes, with the Basal-like (triple-negative breast cancer, TNBC) subtype being the most distinct and the Luminal A, Luminal B, HER2 positive subtypes forming a separate larger group (**Fig. 6c**). We identify intergenic transcriptional markers specific to the Basal-like/TNBC subtype associated with 10 Dual-specificity phosphatases genes (e.g. DUSP1, DUSP5, DUSP7), involved in mitogen-activated protein kinase (MAPK) phosphatase activity. The MAPK cascades are central to cell proliferation and apoptosis, where DUSP1 may contribute to the development of chemoresistance in triple-negative breast cancer^27,28^. TNBC accounts for approximately 15-20% of all breast cancer cases, is most prevalent in women under 40^29^, and presents an aggressive behavior^30^. Similar to breast cancers, the intergenic transcription in Thyroid carcinomas (THCA) allows the detection of different subtypes of thyroid carcinomas (**Fig. 6d**). By using a heatmap representation of the differentially expressed RNAP2 bound regions in Kidney Chromophobe Carcinoma (KICH) samples, we can also observe distinct clusters of up-regulated and down-regulated RNAP2 bound regions, which are potential subtypes of tumours with unique intergenic expression patterns (**Supp. Fig. 14c**). The identification of subtype-specific intergenic transcription sheds light on cancer biology by revealing active regulatory elements and potentially actionable nearby genes with clinical significance.

**Fig 6:**
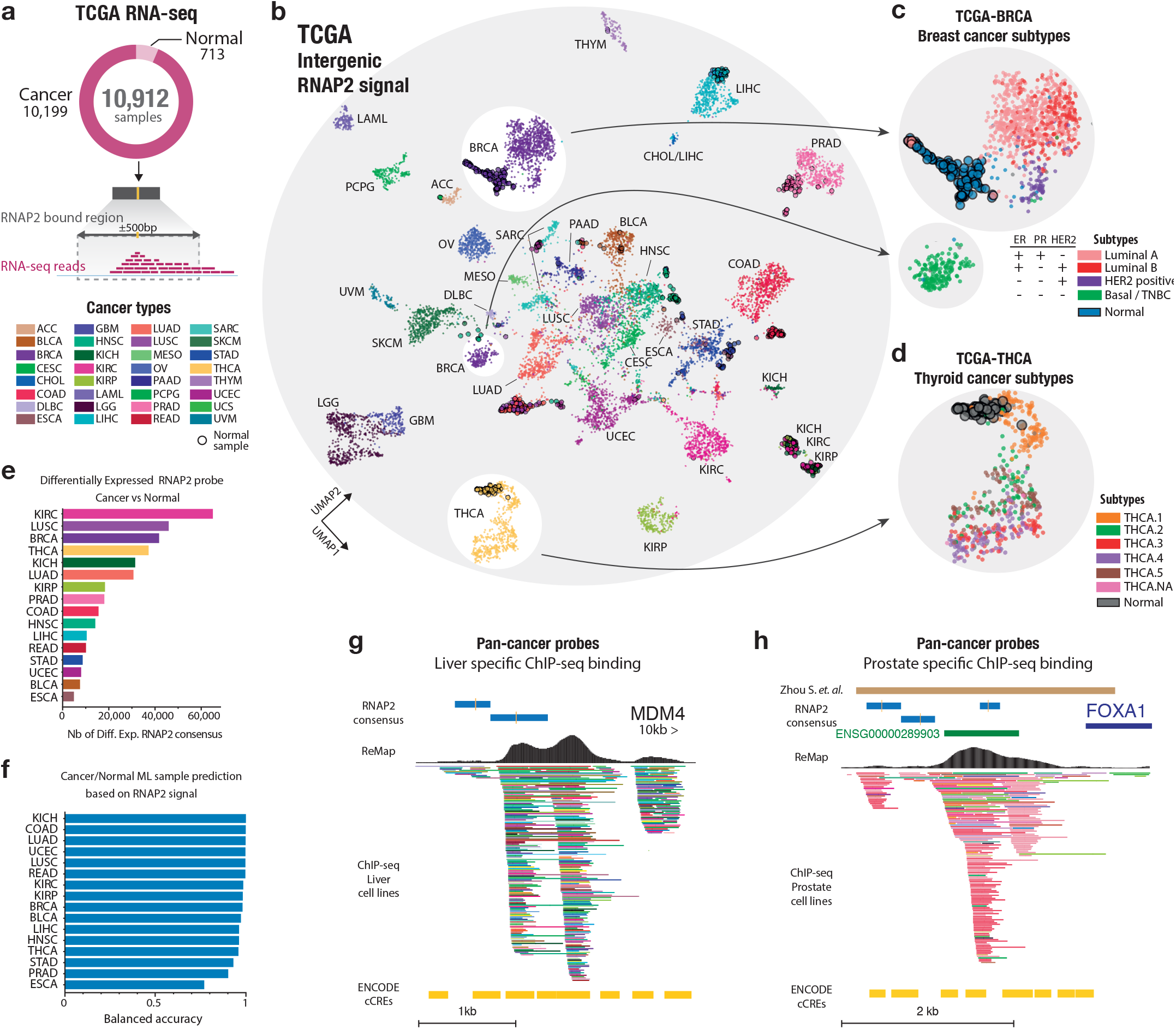
Cancer types and subtypes classification by intergenic transcription at RNAP2 binding sites. **a**, A total of 10,912 TCGA RNA-seq samples were leveraged to capture intergenic signals at standardised RNAP2 1kb bound regions. **b**, A two-dimensional UMAP projection of 10,912 TCGA patients based on intergenic RNAP2 transcriptional signals is shown. Each dot represents a TCGA cancer patient or normal sample, with the colours representing the cancer disease type. White circles highlight Breast cancer (BRCA) and Thyroid carcinoma (THCA) samples. **c**,**d**, Zoomed projections of distinct BRCA and THCA patients (dots) coloured by subtype categories based on intergenic transcriptional signals. Normal samples have larger solid black outlines. **e**, Number of tumour-specific intergenic RNAP2 bound regions differentially expressed in tumours compared to normal samples. **f**. Machine learning classification performance (balanced accuracy) between normal and tumour samples for each cancer type. **g**, Genomic view of a pan-cancer intergenic RNAP2 bound regions differentially expressed in seven or more cancers are shown. Two pan-cancer markers located on enhancers (Enhancer distal, cCREs), near MDM4 gene with ChIP-seq bindings. **h**. Brown bar represent a published cis-regulatory element of FOXA1 harbouring somatic variants in primary prostate tumours^31^. ChIP-seq ReMap tracks are filtered to show TF bindings in Liver or Prostate cell lines specifically.

### Identification of per-cancer and pan-cancer intergenic transcriptional markers

We identified tumour-specific RNAP2 bound regions that were differentially expressed in tumours compared to normal tissues for 16 cancer types, ranging from 65,050 for KIRC (Kidney Renal Clear Cell Carcinoma) to 6,458 for ESCA (Esophageal carcinoma) (**Fig. 6e**). These numbers are consistent with previously identified active enhancers in TCGA cancers^32^. The predictive power of these regions is confirmed as we accurately separate tumours from normal tissues in most cancers using a machine learning classifier (**Fig. 6f, Methods**). To uncover pan-cancer intergenic transcriptional markers that may contribute to tumorigenesis across multiple cancer types, we identified RNAP2 bound regions differentially expressed in a substantial number of cancers (7+ out of 16, **Methods, Supp. Fig. 14b**). We observed a high number of RNAP2 bound regions that do not appear to be differentially expressed in any type of cancer. However, on the other end of the distribution, we observed a significant number of RNAP2 bound regions that are differentially expressed in a greater number of cancers than expected. Specifically, 10,940 RNAP2 bound regions were found to be differentially expressed in more than seven cancers and some in each of the 16 cancers that have corresponding normal tissue samples available. In this set of 10,940 pan-cancer RNAP2 bound regions, we are able to highlight some previously known regions involved in cancer, as well as new loci (**Fig. 6g,h**). For example, we identify two pan-cancer RNAP2 bound regions on enhancers at 10kb upstream of the MDM4 gene, this protein is involved in the repression of the tumour-suppressor TP53 and is a potential therapeutic target in Liver cancer^33^, in Lymphomas^34^ and is overall a target in anticancer therapy^35^. We also highlight a group of pan-cancer RNAP2 bound regions overlapping a known, frequently mutated^31^ FOXA1 enhancer region involved in the proliferation of prostate cancer cells. This region has been identified as one of six cis-regulatory elements in the FOXA1 regulatory plexus harbouring somatic single-nucleotide variants in primary prostate tumors^31^. Our analysis revealed differentially expressed intergenic markers in tumours or tumours subtypes against normal tissues, which may be implied directly or indirectly in the tumorigenesis of tissues. By identifying potential intergenic transcriptional markers, our findings could pave the way for novel therapeutic strategies that target clinically actionable genes.

### Intergenic transcriptional markers showing clinical relevance in cancer

To study the clinical relevance of intergenic transcriptional markers, we investigated the relationship between RNAP2 bound regions expression and overall survival per-cancer and pan-cancer using a Cox Proportional Hazard model (**Supp. Material**, per-and pan-cancer marker lists and count tables available at **Zenodo**^18^). At the per-cancer level, our results showed a smaller number of RNAP2 bound regions associated with overall survival compared to previous differentially expressed RNAP2 expression analyses, with the most being found in Low Grade Glioma (n=18,380), and in average 2,002 per cancer (**Supp. Fig. 15a**). A set of 145 RNAP2 bound regions were found to be associated with overall survival in five or more types of cancer, with most of these bound regions showing a positive association between over-expression and poor survival (Hazard Ratio > 1, **Fig. 7a**). These RNAP2 bound regions were found to be located near genes that are associated with cell cycle, DNA metabolism, repair, and muscle development (**Supp. Fig. 15c**). Perturbation and acceleration of the cell cycle is one of the hallmarks of cancer, and plays a role in tumour progression and prognosis. As examples, we highlight two OS-associated RNAP2 bound regions that are located near known cancer-associated genes and candidate regulatory elements (**Fig. 7b,c**). The first one is located between two non-coding LincRNAs (LINC02873 and ENSG00000288013) at 21 and 33kb, respectively, and is closest to the coding gene SNX19, a sorting nexin, at 128kb. SNX19 has not been clearly linked to cancer survival, but other SNXs have been found to have potential prognostic value in cancers^36^. Decreased expression of SNX1 has been associated with overall survival in colorectal cancer^37^, and down-regulation of SNX2 leads to drug resistance in lung cancer^38^. The RNAP2 pan-cancer bound region reveals a previously unknown correlation between specific survival and expression in cervical, uterine, and breast cancers (**Fig. 7d**). The second RNAP2 bound region is located between the genes TLK1 and METTL8 at 40 and 44kb, respectively (**Fig. 7c**). TLK1 has been linked to poor patient outcomes in multiple types of cancer, including Glioblastoma Multiforme^39^ and Prostate cancer metastasis^40,41^, and is required for DNA replication and chromatin assembly^39^. METTL8 has been identified as a potential biomarker in Hepatocellular carcinoma^42^, and high levels have been linked to patient survival in Pancreatic cancer^43^. We observed that high expression of the pan-cancer RNAP2 bound region depicted in figure 7c is strongly linked to survival in Leukemia, Kidney (KIRP, KIRC), and Lung cancers (**Fig. 7e**). Taken together, these analyses suggest that these transcribed RNAP2 regions, that are mostly unreferenced and undetected, may have a clinically relevant role in cancer and could be useful as potential markers for overall survival. Further studies are needed to fully understand the potential clinical implications of these observations.

**Fig 7:**
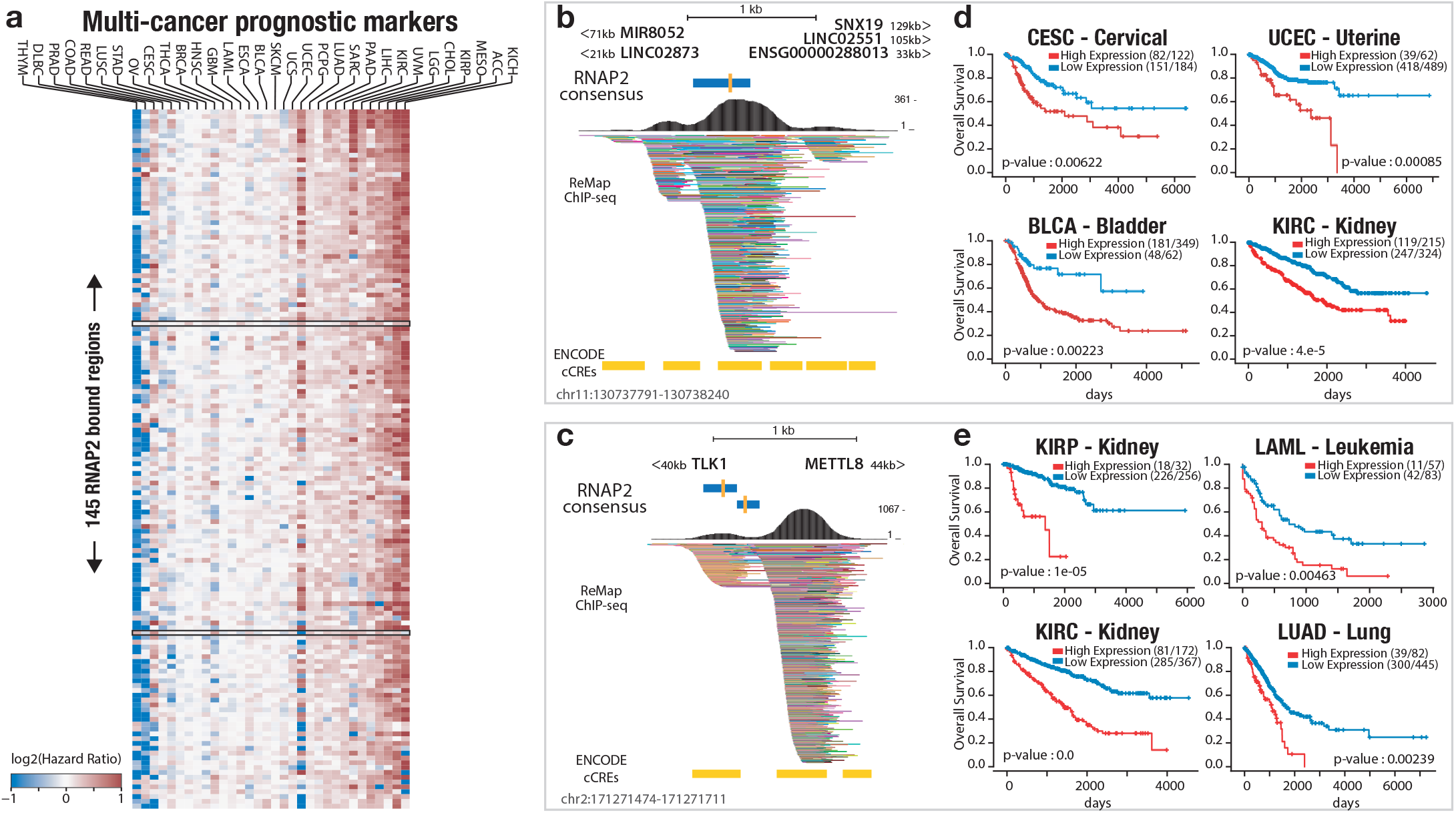
Intergenic transcriptional markers showing clinical relevance in cancer. **a**, Heatmap of 145 transcribed RNAP2 bound regions identified as prognostic markers in multiple cancers. Colour scale depicts log2(hazard ratios) of strong expression associated with good (blue) or bad prognosis (red). Black rectangles highlight two intergenic prognostic markers (RNAP2 bound regions in b-c). **b**, Genomic landscape of an identified multi-cancer prognostic marker blue bar at chr11:130737791-130738240, located upstream (21kb min) and downstream (33kb min) of non-coding and coding genes. Yellow bars indicate candidate Cis Regulatory Elements (cCREs, Enhancer distal) and ChIP-seq binding from ReMap. **c**, Genomic view of the multi-cancer prognostic marker (chr2:171271474-171271711) located 40kb of TLK1 and 44kb of METTL8. d, Kaplan-Meier survival analysis of Cervical, Uterine, Bladder and Kidney TCGA cancer patients with high (red) and low (blue) expression from the intergenic RNAP2 bound region in b. e, Kaplan-Meier survival analysis of Kidney (papillary and clear), Leukemia, Lung cancer patients with high (red) and low (blue) expression from the intergenic RNAP2 bound region in c.

## Discussion

We constructed an atlas of intergenic transcription at RNA polymerase 2 binding sites to connect genomic, transcriptomic and clinical data across normal tissues and cancer samples through the use of a normalised vocabulary for cell lines and tissue types. Our integration framework uses a compendium covering 906 publicly available RNAP2 ChIP-seq profiles allowing to massively probe intergenic transcription across 28,000 expression samples. Our atlas is an efficient way to explore the tissue specificity and activity of core regulatory elements in a tissue. The meta-clustering approach reveals that transcription of intergenic regions is shared across similar tissues and across multiple independent resources. We identified per-cancer and pan-cancer intergenic transcriptional markers associated with known cancer genes, and prognostic intergenic markers predicting the overall survival of patients. Finally, we discover that intergenic transcriptional markers are sufficient to discriminate between subtypes of breast and thyroid cancers.

Our mapping of intergenic transcription stands out from prior efforts to characterise enhancer activities by directly targeting the RNAP2 transcriptional machinery. Traditionally, studies have identified non-coding elements using a single resource such as histone signatures from ENCODE or CAGE transcripts from FANTOM. Our study, however, demonstrates the effectiveness of robust data integration of diverse public RNAP2 ChIP-seq datasets, offering a coherent method to characterise intergenic transcriptional activity in both normal and cancer tissues.

To detect non-coding transcription previous investigations^3,8,23,44^ captured nascent transcripts from GRO-seq or its derivatives. Despite their potential, the limitations of these techniques are twofold. First, the stability and abundance of RNA sources lead to incomplete coverage of the transcribed intergenic genome. Second the limited number of assays result in an underrepresentation of normal tissues and cancer types. We address this by integrating GTEx, TCGA, ENCODE expression data, and provide new perspectives for understanding intergenic activity to track non-coding regulatory circuits across cell lines, normal tissues and cancer types. Most notably, by using signals from TCGA transcriptomes we have established differentially expressed RNAP2 intergenic regions and molecular subtypes of both Breast (eg. TNBC, Luminal) and Thyroid cancers (eg. THC.1). In TNBC, we show that certain RNAP2 DE regions are in the vicinity of DUSP genes that are part of the MAPK signalling pathway. This pathway plays a key role in regulating cell proliferation and apoptosis. DUSP1, in particular, may play a role in the development of chemoresistance in TNBC^27,28^. While we have shown that RNAP2 consensus targets intergenic enhancer elements or proximal enhancers upstream of genes, we also observed RNAP2 consensus located downstream of gene transcription termination sites. Future investigations may allow us to identify potential sites for transcription termination sites (TTS) across our biotype panel.

Transcription of non-coding regions is a fundamental characteristic that is now captured by our RNAP2 intergenic map across cell lines, normal tissues and cancer samples, significantly expanding the analysis horizon beyond gene centric annotations. Our integration framework symbolises the evolution from exploratory studies centred around uncovering new regulatory elements, to a map-focused phase that prioritises the identification of active transcribed elements within specific biological contexts. The significance of our study lies in its ability to enhance our knowledge of activity of non-coding regions in cancer biology and the development of the disease, which may direct us towards therapeutic approaches, and ultimately bring about solutions that could lead to improved patient outcomes.

## Data availability

RNA Pol2 ChIP-seq data are publicly available in NCBI-GEO, and data accessions for ChIP-seq are listed in Supplementary Table 1. The GTEx^15^ eQTL data were obtained from GTEx v8. Human regulatory TF catalogue was obtained from ReMap 2022 release^25^. ENCODE RNA-seq raw sequencing data are available at https://www.encodeproject.org/. TCGA and GTEx RNA-seq raw sequencing data are available under controlled access to ensure appropriate data usage. Access to these protected data must be requested through the dbGaP portal. The Cancer Genome Atlas^16^ (TCGA) RNA-seq BAM files are accessible through dbGaP under accession no. phs000178.v11.p8.c1 (TCGA) and at NCI’s Genomic Data Commons (http://gdc.cancer.gov) under project TCGA. Genotype-Tissue Expression (GTEx) RNA-seq BAM files are accessible through <u>dbGaP</u> under accession no. phs000424.v8.p2.c1 (GTEx) and at the GTEx portal (https://gtexportal.org/home/).

## Code availability

We deposited codes and bioinformatics environments in GitHub at https://github.com/benoitballester/Pol2Atlas. Processed data matrices and files are available at Zenodo^18^. Both data and codes are publicly available for the replication of the whole study.

## Acknowledgements

The authors would like to thank Dr Paul Flicek for constructive criticism of the manuscript, and Dr Sabrina Baaklini for fruitful discussions on meta-clustering. We thank Dr Jeremy J. Day (UAB Heersink School of Medicine) for granting us permission to reproduce the DNA schema on Fig. 1a. This work was supported with ; PhD Fellowship to P.D.L. from the French Ministry of Higher Education and Research (MESR); PhD Fellowship to F.H. from the Provence-Alpes-Côte d’Azur Regional Council (Région SUD); Institut National de la Santé et de la Recherche Médicale (INSERM); The Core Cluster of the Institut Français de Bioinformatique (IFB) (ANR-11-INBS-0013) and the Centre de Calcul Intensif d’Aix-Marseille for granting access to its high performance computing resources. The results shown here are based upon data generated by the TCGA Research Network, the GTEx project, the ENCODE Consortium, the ENCODE production laboratories, and independent laboratories who followed the Open Science principles and submitted raw ChIP-seq data into repositories.

## Contributions

BB initiated, coordinated and supervised the project. PDL manually curated the RNAP2 datasets, EL assisted with primary data curation. FH performed the ChIP-seq reprocessing. PDL performed computational method development and analysed the data, under supervision from LS and BB. PDL and BB generated figures. PDL and BB wrote the manuscript.

## Online Methods

### RNAP2 ChIP-seq data processing

We recovered from NCBI-GEO all existing RNA Pol II (RNAP2) ChIP-seq experiments targeting the POLR2A subunit (n=1,135) in human, following the ReMap procedures and pipeline^25^. Briefly, we manually annotated and standardised the cell line and tissue of origin names (**Supp. Table 1**). Every experiment was downloaded and processed uniformly starting from the fastq files, to quality checks, up to the peak calling stage using the ReMap pipeline. In order to study only the intergenic part of the genome, we filtered out peaks overlapping GENCODE^45^ v38 transcripts ±1kb. We also excluded ENCODE blacklisted regions^17^. We retained peaks with a MACS2 q-value under 10^−5^, and removed uninformative datasets with less than 100 intergenic peaks. In the end, we conserved 906 out of 1,135 datasets after all Quality Checks (**Supp. Fig. 2a**,**b**.).

### High level biosample annotation

Due to the very large biological diversity of the experiments, it is necessary to have a high level annotation to make the interpretation of the results easier, as well as comparing results between datasets. We annotated samples according to their tissue of origin, with the simplified GTEx tissue (30 tissues) annotation as a baseline, to which we added additional tissues : bone, eye, embryo and trachea. To make some results more interpretable, we grouped similar tissues (eg. various brain tissues into ‘Brain’) obtaining an annotation with 18 categories (**Supp. Table 3**). A full sample-annotation table is available in **Supp. Table 1**.

### Construction of the intergenic RNAP2 Atlas

A naive approach to delineate groups of RNAP2 peaks corresponding to a similar biological signal across experiments would be to merge overlapping peaks. However, when the number of experiments is large, the entire genome becomes covered with peaks which makes this approach impractical. To create consensus RNAP2 peaks, we first computed the density function of the peak summits (the single base pair genomic location with the maximum signal of the peak) across each chromosome. Due to the inherent inaccuracy on the summit position of these sequencing techniques and the undersampling, this estimate is extremely noisy. To reduce the amount of noise, we applied a Gaussian filter to this density function across the genome (Fig. 1a). Consensus peaks were defined at each local minima of the smoothed density function. A peak belongs to a consensus if its peak summit falls in between the identified flanking local minimas. The boundaries of the defined consensus peak were reduced to the ones of the farthests peaks. When processing stranded peak data (for example CAGE or RAMPAGE), these steps were run once for each strand. By default 1/8th of the average peak size was used as the standard deviation of the Gaussian kernel, and to be valid each consensus was required to contain at least 2 peaks from different experiments. Consensus peaks centroid were defined as the mean position of the peak summits. The middle of the peak was used, if a summit coordinate was not available. A binary data matrix was generated to summarise all datasets. For each consensus peak, this matrix stores if a biosample has a RNAP2 peak that belongs to it, similar to the DNAse 1 binary matrix from ENCODE^22^. A schematic of the whole approach is available in **Supp. Fig. 1**.

### RNAP2 atlas visualisation and clustering

To visualise the similarity between datasets, we applied UMAP with the Yule similarity, with 30 neighbours and the minimum distance set to 0.5. To visualise the similarity between RNAP2 consensuses, we use the Sorensen-Dice similarity, 30 neighbours and the minimum distance is set to Other parameters were left to default. For the consensus peaks UMAP, to highlight consensus peaks specific to a biotype annotation, each consensus peak was coloured by its most frequent biotype annotation. To do so, we compute the sum of the number of peaks per dataset of each annotation at each consensus (*s*_*i,j*_), which is then normalised by the total number of peaks for each annotation 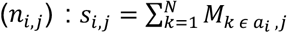 and 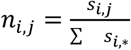, where *a*_*i*_ is a set storing the index of each datasets belonging to annotation i, M is the dataset binary matrix, N the number of experiments, and j the consensus index. This prevents over-represented annotations or annotations with some datasets with a large number of peaks to annotate most of the consensus peaks. We chose as representative for consensus j the annotation i for which *n*_*i,j*_ is the largest.

Finally, to identify consensus peaks that are not condition specific, each one is linearly grayed according to its Gini-Simpson index : 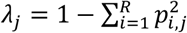, where R is the number of annotations, and 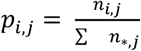. The Gini-Simpson index is a measure of diversity : in this study, it tends towards one if the annotations are equidistributed, and is equal to zero if the consensus only has peaks belonging to datasets with the same annotation.

A three steps Hierarchical Clustering (HC) approach was used to order datasets and RNAP2 consensus peaks. First, we performed a UMAP dimensionality reduction to 10 dimensions, and used the same metric as the 2D UMAP transform. This step allows the use of any metric, as UMAP optimises to a lower dimensional space using the euclidean distance, which is used by k-means and Ward HC, and also improves k-means and HC quality. Second, we reduced the effective number of points using k-means clustering and grouped very similar points into 50,000 clusters (step performed only when >50,000 points). This approach is documented and allows to scale Ward HC to very large datasets^46^. Third, we performed Ward HC on the k-means clusters centroids using the fastcluster library. The bottom part of the heatmap displays each *p*_*i,j*_ as defined in the previous section as a stacked barplot. A Shared Nearest Neighbour (SNN) Graph Clustering approach was used to identify clusters of RNAP2 consensus peaks. This approach is common in single-cell RNA sequencing (scRNA-seq) analyses to identify clusters of cells without a priori on the number of clusters. To scale to a large number of points to cluster, we used an Approximate Nearest Neighbour (ANN) method to build the NN graph (python library pynndescent^47^). This approach avoids the quadratic time complexity of building exact nearest neighbours, can use any metric and runs in an almost linear time complexity. The Sorensen-Dice coefficients were used to measure distances between points. In the SNN graph, vertices are weighted by the number of shared nearest neighbours between the two nodes. To identify communities in the SNN graph, we used the Leiden graph clustering algorithm implemented in the python leidenalg^48^ library.

### Extending the integrative approach to H3K27Ac ChIP-seq

We collected all H3K27Ac ChIP-seq experiments from ENCODE, retrieved processed files in bed narrowPeak format, mapped for hg38, and without audit error appearing on the sample metadata (n=890 samples). Each sample was annotated with the same biotype methodology as our RNAP2 consensus. The same integrative approach and settings as the RNAP2 atlas presented above were run on this dataset.

### Comparison with reference databases from other large-scale efforts

The RNAP2 atlas was intersected against GENCODE^45^ v38, LNCipedia^49^ v5, FANTOM5^7,50^, ReMap^25^ 2022 and Repeat elements downloaded from UCSC^51^ (hg38). Intersections are computed using the centroid of RNAP2 consensus (1bp) against the whole genomic features. The PyRanges python library was used to compute intersections between genomic features. We computed overlap enrichments for the whole dataset using a binomial test where : n, the number of trials, is the number of RNAP2 consensus; p, the probability of intersection, is the base pair coverage of the feature of interest divided by the the coverage of the intergenic regions (+-1kbp from genes, excluding ENCODE blacklisted regions); k, the observed number of successes, is the number of RNAP2 consensus intersecting the feature of interest; The fold change is computed as 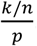. We computed overlap enrichments for subsets of the whole RNAP2 atlas using a hypergeometric test, which removes the RNAP2-specific intersection bias : where N, the population size, is the number of RNAP2 consensus; K, the number of successes in the population, is the number of RNAP2 consensus intersecting the feature of interest; n, the number of draws, is the number of RNAP2 consensus of the subset of interest; k, the number of observed successes, is the number of RNAP2 consensus of the subset of interest intersecting the feature of interest.

### Epigenetic enrichments

We downloaded every 15 states epigenome available from ROADMAP^52^ (hg38). We intersected (consensus centroid only) each consensus peak with each epigenome to get the epigenetic state of each consensus in each epigenome. We computed the proportion of epigenetic states for the subsets/clusters of RNAP2 consensus of interest for each epigenome (i.e. the sum of the epigenetic states proportions is equal to one in an epigenome). We used the “GROUP” column of the epigenome metadata to annotate and group epigenomes. We used a paired t-test to statistically assess the difference in proportions between subsets of RNAP2 consensus across epigenomes. We downloaded H3K27Ac ChIP-seq and ATAC-seq processed bam files from ENCODE for Heart, Liver and T cells samples (**Supp. Table 2**). We used deeptools to compute the mean profiles at each RNAP2 consensus of the studied clusters (+-5kb from centroid).

### Gene Ontology enrichments of nearby genes

To assign consensus peaks to genes, we used a similar heuristic as GREAT at default settings : a basal domain of 5kb upstream and 1kb downstream, extended in both directions up to 1Mb or the nearest basal domain (whichever is the closest). For each gene we obtained the number of consensus peaks in its regulatory region, for all the consensus peaks (n) and its subset of interest (k). To compute Gene Set enrichments, we used a Negative Binomial GLM : *ln*(*μ*) = *β*_0_ + *β*_1_ × *G* + *E*. Where G is equal to 1 if the studied gene belongs to the Gene Set of interest and 0 otherwise. The term E corrects for the intersection bias of the background regions, with E being the expected number of hits for a particular gene : 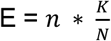, with K being the number of query regions, and N the number of background regions. We tested whether *β*_1_ is greater than zero using a Wald Test. The model is fitted using the python statsmodels^53^ library. We considered GO terms with more than 3 genes and less than 1,000, and applied the Benjamini-Hochberg FDR correction. The approach is similar to Chip-Enrich and Poly-Enrich, which has shown that gene-wise modelling is required to reduce false discoveries, but these two methods do not offer a model for our case, where the query regions are a subset of a set of background regions. To improve the readability of the GO enrichments, we identified clusters of GO terms given to similar genes using a graph clustering approach to reduce term redundancy. Starting from a binary matrix with genes as columns and significant (5% FDR) GO terms as rows, we built a nearest neighbour graph of GO terms using the Yule metric. We performed graph clustering on this NN graph and chose the GO term with the smallest P-Value as the cluster representative for each cluster.

### GWAS traits and summary statistics

We used Stratified LD-Score Regression^54^ (S-LDSR) to compute enrichments of heritability phenotypes for subgroups of consensus peaks. We downloaded all available GWAS summary statistics files from UK Biobank^24,55^ (http://www.nealelab.is/uk-biobank), and only kept traits with strong heritability (noted “z7”, as recommended by the documentation for this kind of analysis). RNAP2 consensus coordinates were lifted to hg19 (UCSC liftover) which caused 873 RNAP2 consensus to be removed. An SNP was assigned to a RNAP2 consensus if it overlaps any part of the RNAP2 consensus. The LDSC pipeline was run at default settings and bonferroni correction was applied on the obtained p-values. Heritability enrichments are defined as : 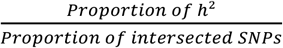, where h^2^ is the SNP-based heritability (see LD-Score paper^54^).

### RNA-seq expression quantification of RNAP2 bound regions

We quantified RNA-seq expression on RNAP2 bound regions using featureCounts^56^ similar to RNA-seq gene quantification. Instead of genes as sampling points, RNAP2 bound regions were used and standardised to 1kb long, centred on the consensus centroids (±500bp). Multi-mapping reads were excluded. Given the similarities of our data with scRNA-seq datasets, we employed several methods commonly used in the scRNA-seq field. Our data has a large number of samples (equivalent to cells) with much lower read counts compared to traditional gene-centric RNA-seq experiments. Before conducting each analysis, we preprocessed the data by filtering out RNAP2 bound regions that did not have at least one read count in three samples. This filtering step helped to remove noise and increase the quality of the data. The “gene-centric” GTEx count table was retrieved from GTEx^15^.

### Count normalisation and transformation

An overview of our count processing is available in **Supp. Fig 6**,**7**. Counts were normalised using the scran pooling and deconvolution^57^ approaches, as RNAP2 bound region counts have a large fraction of zeroes causing issues on approaches such as DESeq2’s median of ratios. A small modification of the method was used to compute the size factors only using the top 5% “most detectable” bound regions. We defined detectability as the number of samples that have at least one read at an RNAP2 bound region. To break ties, we computed detectability at 2 reads, 3 reads… up to 5 reads, which is sufficient to break most ties. This helped to reduce the number of non-expressed RNAP2 bound regions or RNAP2 bound regions that are expressed in a single condition only, which should not be considered for an optimal normalisation. This can be seen as something analogous to the use of the geometric mean in DESeq2’s median of ratios, which considers only genes with at least 1 read in each sample. RNAP2 counts were transformed using the Pearson residuals of a regularised Negative Binomial (NB) model, similarly to what is used in SCTransform^58^. This kind of transform has been shown to better discriminate between biological conditions in scRNA-seq experiments, as well as reducing batch effects caused by differences in sequencing depths at small counts. The model for a gene/RNAP2 bound region expression is : *ln*(*μ*) = *βX* + *ln*(*s*). Where µ are the predicted means for a bound region/gene for each sample, *β* are the fitted model coefficients, *X* is the design matrix, and s are the count normalisation factors for each sample. In this work we use a simple intercept but the model allows more complex experimental designs. We use the following NB(µ, α) variance formulation : *V*(*μ*) = *μ* + *α μ*^2^. We fitted a trendline of the overdispersion parameter *α* as a function of the mean, so we obtain a regularised estimate of the variance that only depends on the mean. To do so, we binned genes/RNAP2 bound regions into 20 groups according to their quantile of mean expression, evaluated each gene/bound region overdispersion parameter, find the modal value of overdispersion within a group using a kernel density estimate (with Silverman’s rule to estimate bandwidth), and linearly interpolate results between each group/quantile of mean expression. We only used up to 5000 RNAP2 bound regions/genes to fit the mean/overdispersion relationship to speed up computations. The python statsmodels library was used to fit the NB models with the more robust Nelder-Mead solver instead of BFGS. The pearson residuals are then computed as following :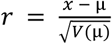, where x is the count value. A custom python implementation was employed as the SCTransform package failed to run on the RNAP2 count matrices, possibly due to much larger counts than UMI scrna-seq experiments, causing numerical instability when fitting the models. The original implementation clips the pearson residuals at 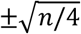 by default, where n is the number of cells/samples, in order to reduce the influence of outliers. We found these bounds to be quite small when dealing with smaller sample sizes, which can remove biological signal. Instead, we clipped values at 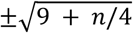, creating larger bounds for small sample sizes without changing the large sample size behaviour.

### Differential expression

To identify differentially expressed (DE) RNAP2 bound regions between tumour and normal tissues, we performed a t-test on the Pearson Residuals with a FDR cutoff of 5%. We constrained DE RNAP2 bound regions to have an absolute log2 Fold Change above 0.25, and to be detectable in at least 10% or 2+ samples (whichever is the largest) of either class (normal/tumour). For the detection of Tumour Subtype specific markers, we compared the expression of samples of a subtype to reference normal samples. We used the same significance cutoffs. We considered a marker to be subtype-specific only if it appeared for this subtype. Linear modelling methods such as DESeq2 ran out of memory on large datasets and required an excessive computation time. A t-test was used to accommodate large datasets (100+ samples x 180 000 bound regions in most datasets) and to keep an uniform processing for each dataset for our cross-dataset analyses. With sufficiently large sample sizes, the t-test yields robust markers, although with less statistical power (**Supp. Fig. 7 c-d**). To evaluate our approach on a much smaller dataset with less statistical power, we selected samples from two similar types of heart tissues from GTEx and downsampled to obtain a n=3 comparison. Here, we used DESeq2 to maximise statistical power. We performed 100 random sampling iterations to obtain 3 samples for each tissue, evaluated DE in each iteration, then kept bound regions supported as DE in at least half of the downsampling iterations. To evaluate the relationship between sample size and statistical power, we performed 10 downsampling iterations for each sample size. GREAT v4.0.4 analyses were performed on BRCA TNBC specific RNAP2 bound regions, identifying enrichment of the “MAP kinase tyrosine/serine/threonine phosphatase activity” GO:0017017 term (Binom FDR Q-Val 1.34e-14, with TNBC specific RNAP2 bound regions associated with 10 DUSPs genes (recherche.data.gouv.fr).

### Survival analysis

For survival analysis, we fit a linear Cox Proportional Hazards regression model on the Pearson residuals using the Python Lifelines library, and use a 5% FDR threshold. Kaplan-meier survival curves were created using the kaplanmeier python library. The maxstat R library was used to obtain the optimal expression cutpoint as well as the associated maximally selected logrank statistic p-value (using the most accurate “condMC”‘ method with 100 000 samples to compute the p-value).

### Identification of pan-cancer markers

To identify RNAP2 bound regions whose expression is associated with survival or DE (separately) in tumour tissues in multiple cancers (“pan-cancer markers”), we randomly selected for each cancer the same number of marker bound regions as observed in this cancer. This process was repeated 100 times to obtain the expected distribution of the number of cancers in which a bound region is a marker. We identified the “pan-cancer threshold” as the threshold where less than 5% of the observed markers are expected to belong to the null distribution (equivalent to 5% FDR, see Supp. Fig. 13b). This approach allowed us to set a statistically meaningful threshold to identify bound region that are markers in more cancers than expected at random, instead of an arbitrary threshold.

### Unsupervised feature selection, dimensionality reduction and predictive models

Feature selection in scRNA-seq is a common step that allows to remove a large fraction of potentially uninformative bound regions/genes (ie those with very low expression or those with ubiquitous expression, which are not informative of the sample/cell biology). Typically, around 2000 to 3000 genes are kept in scRNA-seq experiments, but this number is generally tuned for each experiment. To automatically select “highly variable” features for each dataset, we computed the sum of the squared pearson residuals, which are asymptotically following a *χ*^2^distribution with n -p degrees of freedom, n being the number of samples, and p the number of parameters of the model (1 in our case). We performed an upper tail test for each gene/bound region and kept bound regions at a FDR of 5%. This selects only sufficiently expressed genes above the mean-variance trendline, and due to the clipping of the pearson residuals also removes outliers with an extreme variance (**Supp. Fig. 7a**). We performed PCA on the Pearson Residuals of these highly variable features. To automatically identify the optimal number of Principal Components, we used Horn’s Permutation Parallel Analysis, which has been found to be one of the most effective approaches to identify the number of components in factor analysis (cit). This approach generates row–wise permutations for each feature, computes PCA on these permuted datasets, then the selected number of components is the threshold at which the eigenvalues from the randomised dataset are larger than the real dataset. We performed 3 permutations due to the computational cost of this approach, which is acceptable as the randomised eigenvalues are very stable on large matrices (**Supp. Fig. 7b**). We used the fast “randomised” solver from the python sklearn^59^ library to compute PCAs.

For UMAP visualisation, we used 30 neighbours, a min_dist parameter of 0.5, Pearson correlation as the metric and use data in PCA space as input. For heatmaps, we used a similar approach as the RNAP2 heatmap, except that the data was used in PCA space as input to the UMAP pass for the samples, and used Pearson correlation as the metric for both samples and RNAP2 bound regions. The predictive model uses a Catboost gradient boosted decision tree model that takes as input the data in PCA space. Default settings were used with the exception of balanced class weights (where each sample is reweighted by class proportion). We used balanced accuracy (where each sample is reweighted by class proportion) as the main metric to evaluate the model over a stratified 10-Fold Cross Validation.

### Identification of per tissue markers and “meta-clustering”

For each dataset (TCGA, ENCODE, GTEx, RNAP2), we identified markers for each annotation (ie Pol2+Liver, GTEx+Blood). To identify markers in the three RNA-seq datasets, we performed a group-versus-rest, one sided t-test on the Pearson Residuals. We kept over-expressed bound regions with log2 Fold Change above 0.25, and detectable in at least 10% or 2+ samples (whichever is the largest). For the RNAP2 dataset, we performed an hypergeometric test for each RNAP2 consensus, where : N, the population size, is the number of peaks across all experiments; K, the number of successes in the population, is the number of peaks across all experiments with the annotation of interest; n, the number of draws, is the number of experiments that has a peak at the studied consensus peak; k, the number of observed successes, is the number of experiments with the annotation of interest that has a peak at the studied consensus peak. We used a BH FDR cutoff of 5% in both cases. This yields a binary vector which indicates whether a RNAP2 bound region is a marker or not for each Dataset+Annotation. We removed RNAP2 bound regions which are markers in more than 10% of the dataset+annotation combinations or in less than two dataset+annotation. An Average Linkage clustering using the Yule binary metric was performed, which we found to be less sensitive to the number of identified markers.

### Tissue specific eQTL enrichments

We downloaded per tissue eQTL data from GTEx “GTEx_Analysis_v8_eQTL.tar” and used the list of significant eQTL-gene pairs for each tissue. For each SNP listed as an eQTL we stored whether it is listed as an eQTL or not in each tissue. SNPs listed as eQTLs in more than 10% of the tissues (6 or more) were removed to keep variants that are likely located in tissue specific regulatory regions. Using the list of per dataset, per tissue marker RNAP2 bound regions, we kept RNAP2 bound regions that are markers in less than 10% of the tissues in a dataset (6 or less) and removed non-marker RNAP2 bound regions. We removed tissues having less than 50 markers left after this step. We computed pairwise intersection enrichment p-values between the tissue-specific eQTLs and the marker RNAP2 bound regions (SNP intersection against whole bound region). We computed an hypergeometric enrichment p-value for each of these intersections as following : N, the population size, is the number of RNAP2 bound regions (after filtering); K, the number of successes in the population, is the number of RNAP2 bound regions intersecting an eQTL from the eQTL-wise tissue of interest; n, the number of draws, is the number of marker RNAP2 bound regions of the second tissue of interest; k, the number of observed successes, is the number of marker RNAP2 bound regions of the second tissue of interest intersecting an eQTL from the eQTL-wise tissue of interest.

