## Supplementary_materials-and-methods for "Normal and cancer tissues are accurately characterised by intergenic transcription at RNA polymerase 2 binding sites"

**Supplementary figures** **2**
Figure S1 : Large scale integration methodology of RNAP2 ChIP-seq 2
Figure S2 : Characteristics of the RNAP2 atlas. 3
Figure S3 : Genome-wide large scale integration of 890 Human H3K27Ac Histone ChIP-seq experiments. 4
Figure S4 : Tissue-specific epigenetic states of RNAP2 consensuses. 5
Figure S5 : Tissue-specific biological characteristics of RNAP2 consensuses. 6
Figure S6 : Flowchart of the RNA-seq pipeline 7
Figure S7 : Mean-variance trendline, Feature selection and PCA Permutation Parallel analysis 8
Figure S8 : RNAP2-bound regions captures a majority of intergenic transcriptional signal 9
Figure S9 : Intergenic transcription by itself is sufficient to characterise biological conditions 10
Figure S10 : Tissue-specific regulatory variants are enriched within tissue-specific Intergenic transcripts 11
Figure S11 : The intergenic transcriptional signal is not driven by end-of-gene transcription 13
Figure S12 : Differentially expressed RNAP2-bound regions can be detected at smaller sample sizes. 14
Figure S13 : Per-biotype robustly over-expressed markers display meaningful disease-associated heritability enrichments 16
Figure S14 : Non coding transcription captured at RNAP2-bound regions discriminates normal and tumour tissues. 18
Figure S15 : Non coding transcription captured at RNAP2-bound regions is prognostic of the patient's survival. 19

### Supplementary figures

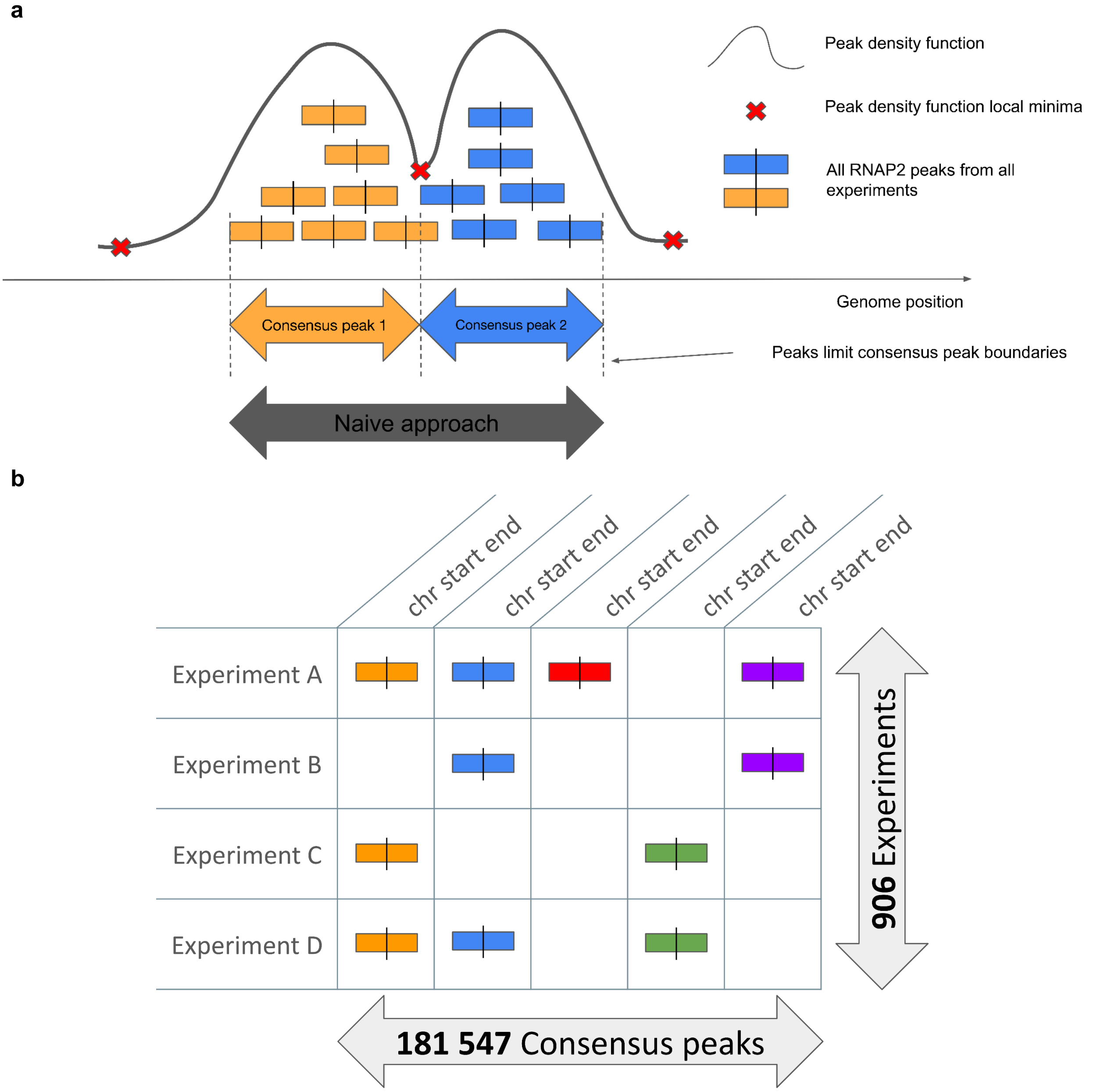


**Figure S1 : Large scale integration methodology of RNAP2 ChIP-seq**

Schematic of our post peak-calling integration methodology : **a.** Identification of consensus peaks via a peak density-based approach. Naive approach refers to a simple merge on overlap. **b.** Summarization of all datasets / consensus peaks in a binary matrix storing the presence or absence of RNAP2 at each consensus peak in each experiment.

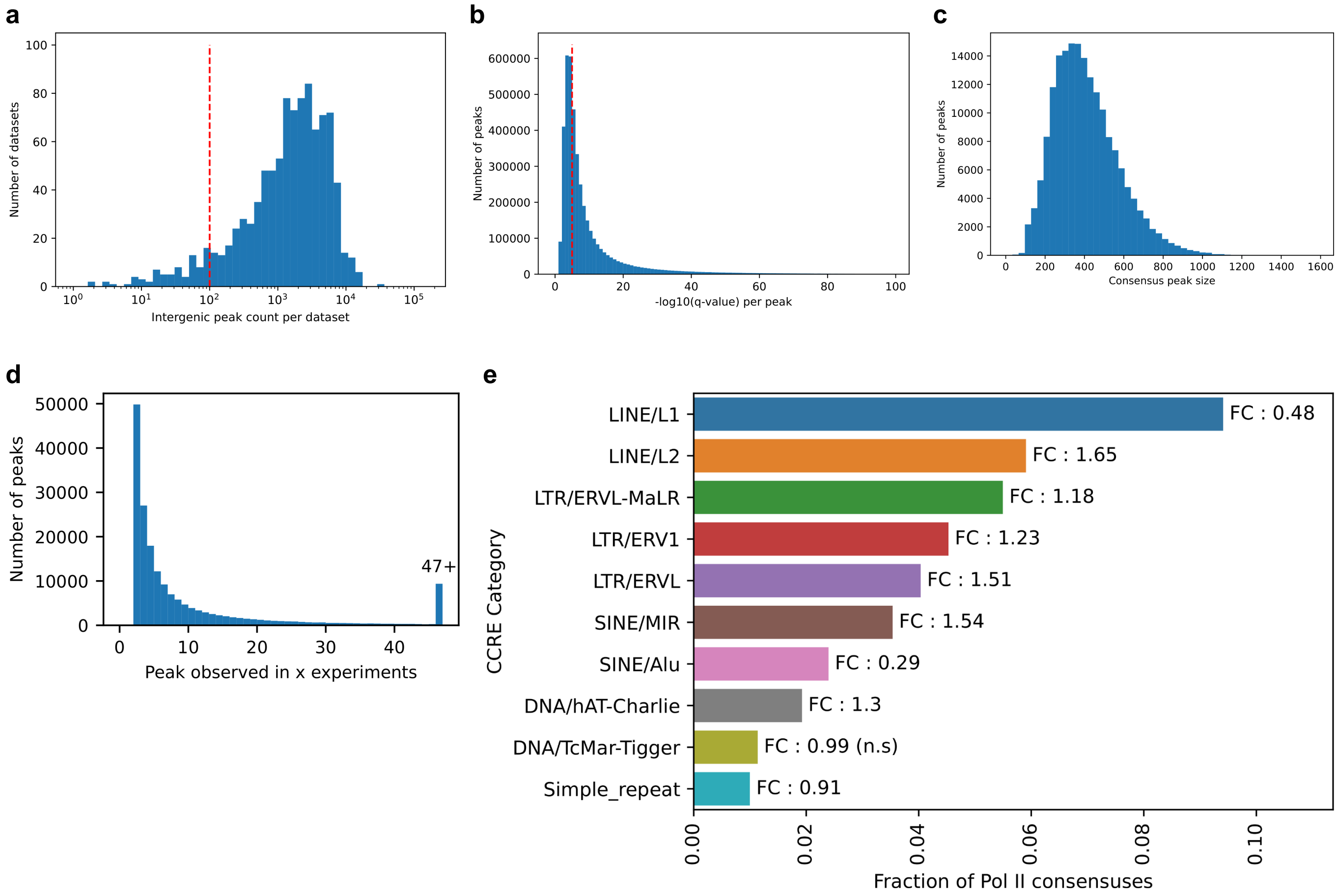


**Figure S2 : Characteristics of the RNAP2 atlas.**

**a.** Histogram of the distribution of the number of Intergenic RNAP2 peaks per dataset. Red line indicates minimal cutoff for a dataset to be retained. **b.** Histogram of the distribution of intergenic peak q-value (all experiments). Red line indicates minimal cutoff for a peak to be retained. **c.** Histogram of the distribution of the intergenic consensus peak sizes. **d.** Distribution of the number of peaks contributing to an intergenic consensus peak. **e.** Fraction of consensus peak intersected for the top ten most intersected repeat families. FC corresponds to fold change enrichment versus random regions (see methods). All results are statistically significant unless otherwise mentioned.

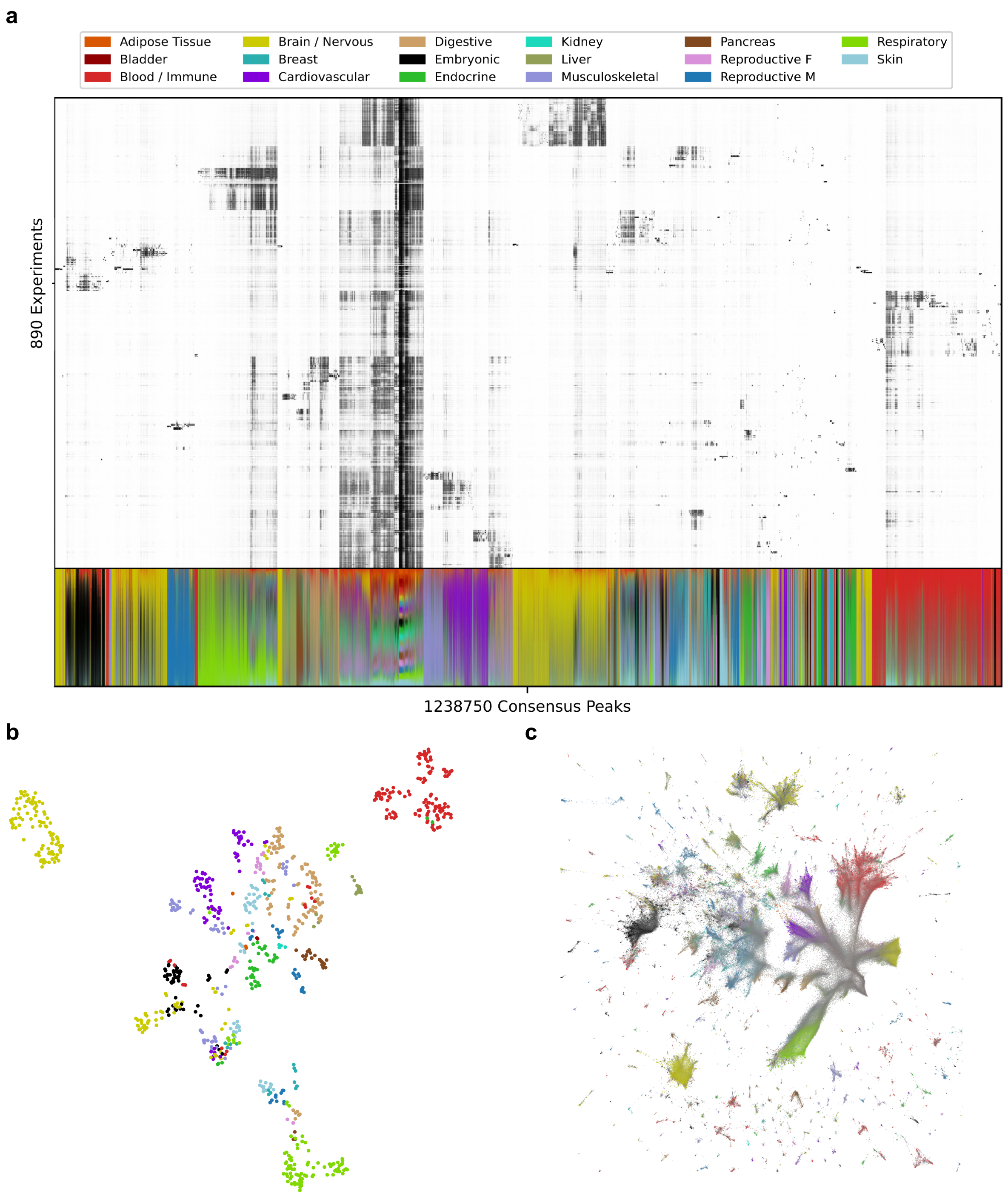


**Figure S3 : Genome-wide large scale integration of 890 Human H3K27Ac Histone ChIP-seq experiments.**

**a.** H3K27Ac occupancy in 1,238,750 consensus regions across 890 biosamples. Lower panel indicates the normalised contribution of a biotype, in terms of peaks, to each consensus. **b.** Two-dimensional Uniform Manifold Approximation and Projection (UMAP) projection of all 890 H3K27Ac ChIP-seq datasets. **c.** UMAP projection of all H3K27Ac consensus according to their binding patterns, coloured by dominant biotype.

**a**

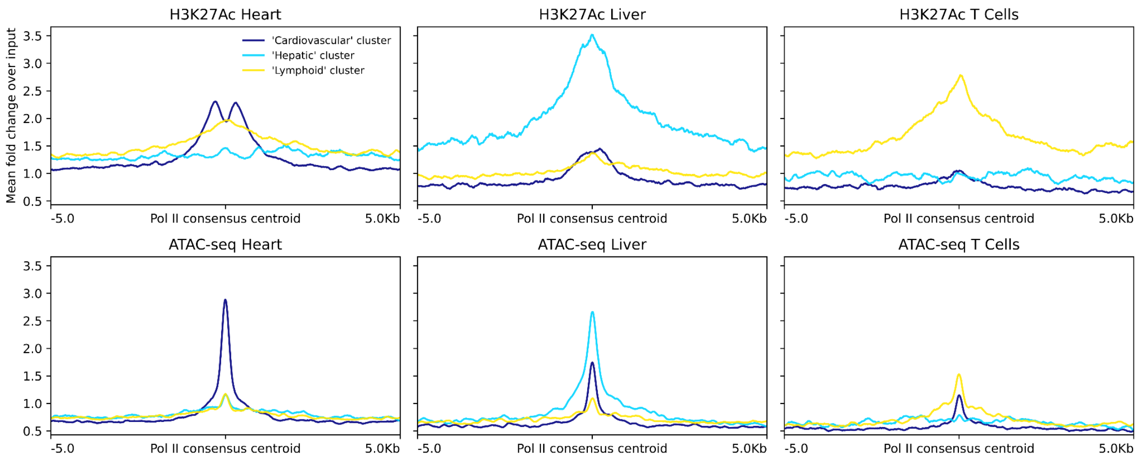


**b**

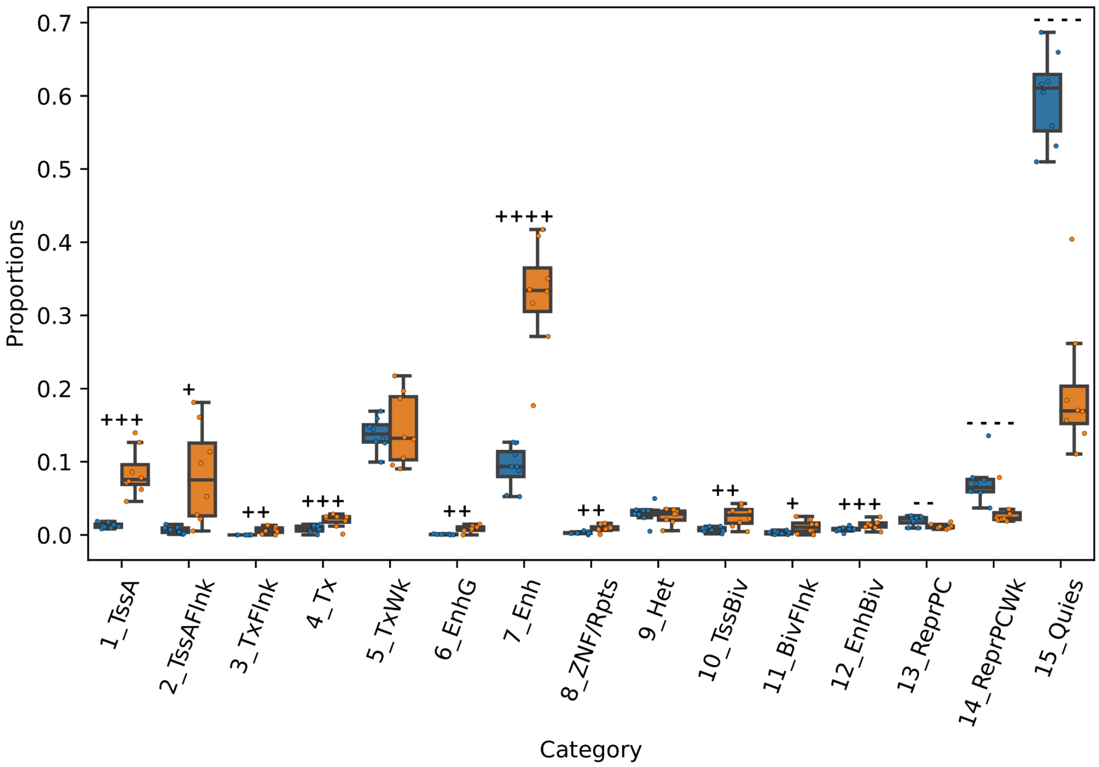


**Figure S4 : Tissue-specific epigenetic states of RNAP2 consensuses.**

**a.** H3K27Ac ChIP-seq and ATAC-seq profiles at clusters of RNAP2 consensuses. Selected clusters are respectively #5, #10 and #20 for “Cardiovascular”, “Hepatic” and “Lymphoid” (largest representative clusters, see supplementary data). **b.** Proportions of ChromHMM epigenetic states in the “Embryonic” cluster against other Intergenic RNAP2 consensuses. (* : $p < 0.05$, ** : $p < 0.01$, *** : $p < 0.001$, **** : $p < 0.0001$; + or - indicate sign of mean difference; two-sided paired t-test).

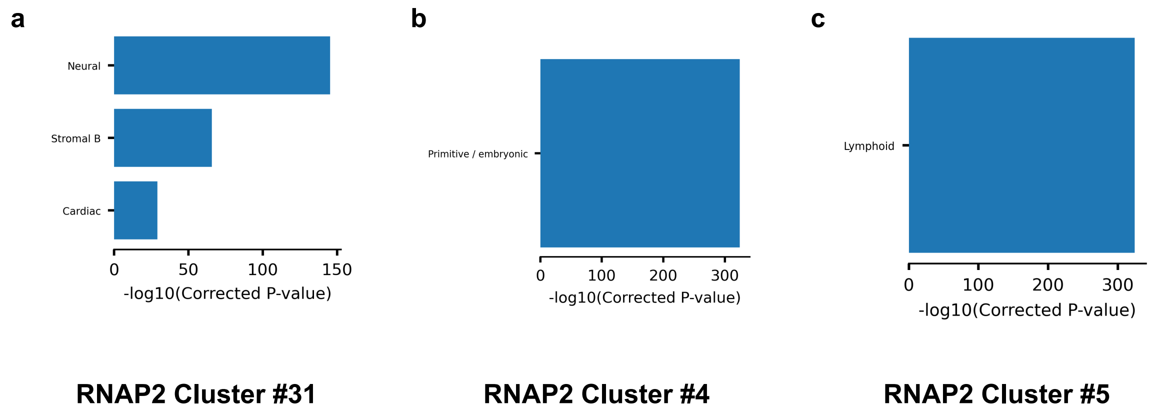


**Figure S5 : Tissue-specific biological characteristics of RNAP2 consensuses.**

Significant enrichment p-values of genomic intersections between DNAse dominant component clusters from ENCODE and **a.** RNAP2 Cluster #31, **b.** RNAP2 Cluster #4, **c.** RNAP2 Cluster #5.

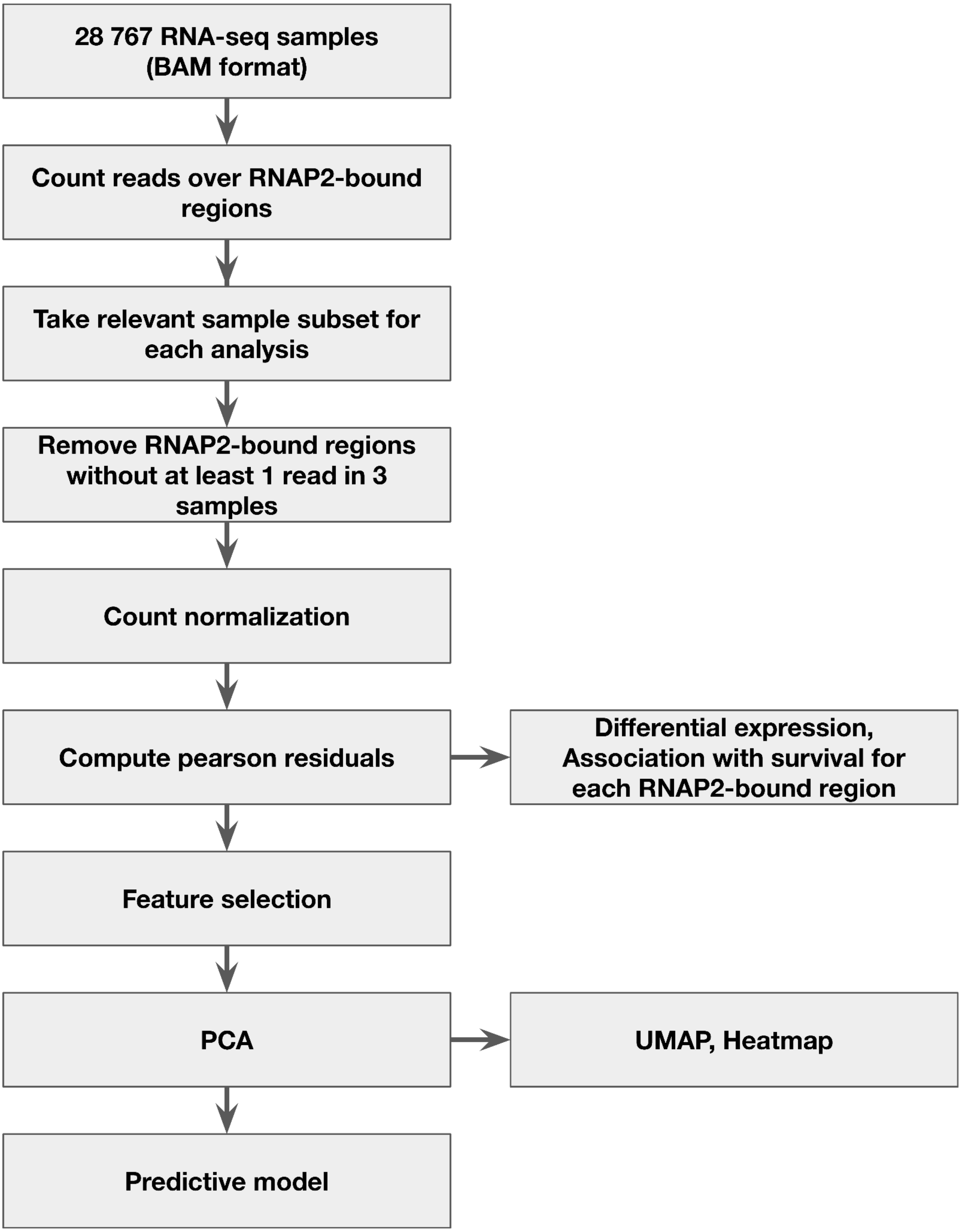


**Figure S6 : Flowchart of the RNA-seq pipeline**

Simplified schematic of the RNA-seq processing pipeline. See methods for additional details.

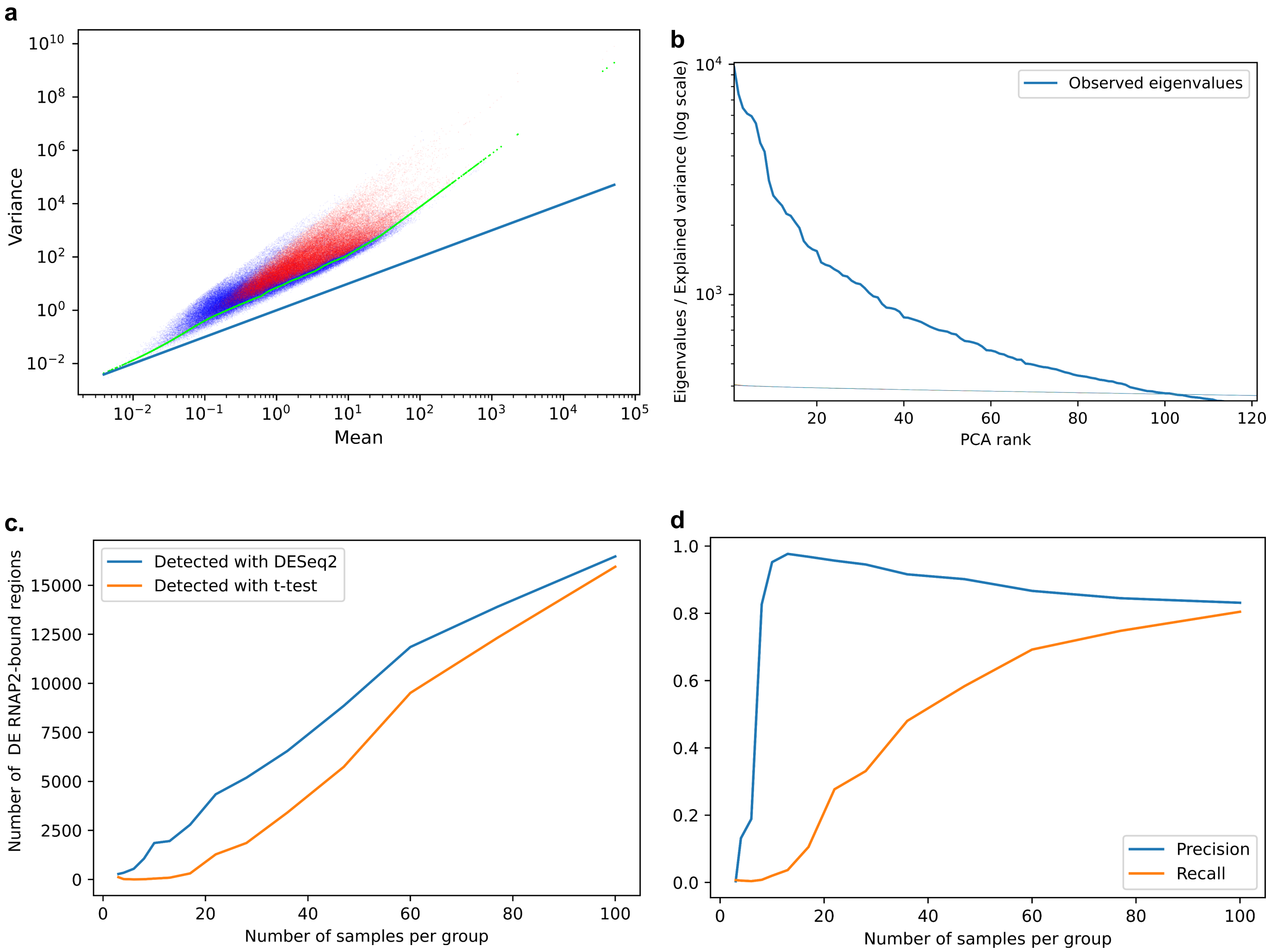


**Figure S7 : Mean-variance trendline, Feature selection and PCA Permutation Parallel analysis**

**a.** Scatterplot of the mean and variance of the normalised counts of the 180 000+ RNAP2-bound regions in the ENCODE rna-seq dataset. Red dots are selected "highly variable" RNAP2-bound regions, green dots represent the fitted mean-variance trendline. Blue line is a Poisson mean-variance relationship. **b.** Observed eigenvalues (or explained variance) for each component of the PCA performed on the pearson residuals of selected RNAP2-bound regions in the ENCODE rna-seq dataset. 100 thinner lines (stacked on the graph) are corresponding to the PCA eigenvalues of each of the 100 permutations of the dataset. Here, only the 102 first components are retained. **c.** Average (over 10 downsampling iterations) number of detected DE RNAP2-bound regions between the two heart tissues in function of the number of samples per group. **d.** Average (over 10 downsampling iterations) precision and recall of the t-test in function of the sample size, using DESeq2 DE-bound regions as a reference.

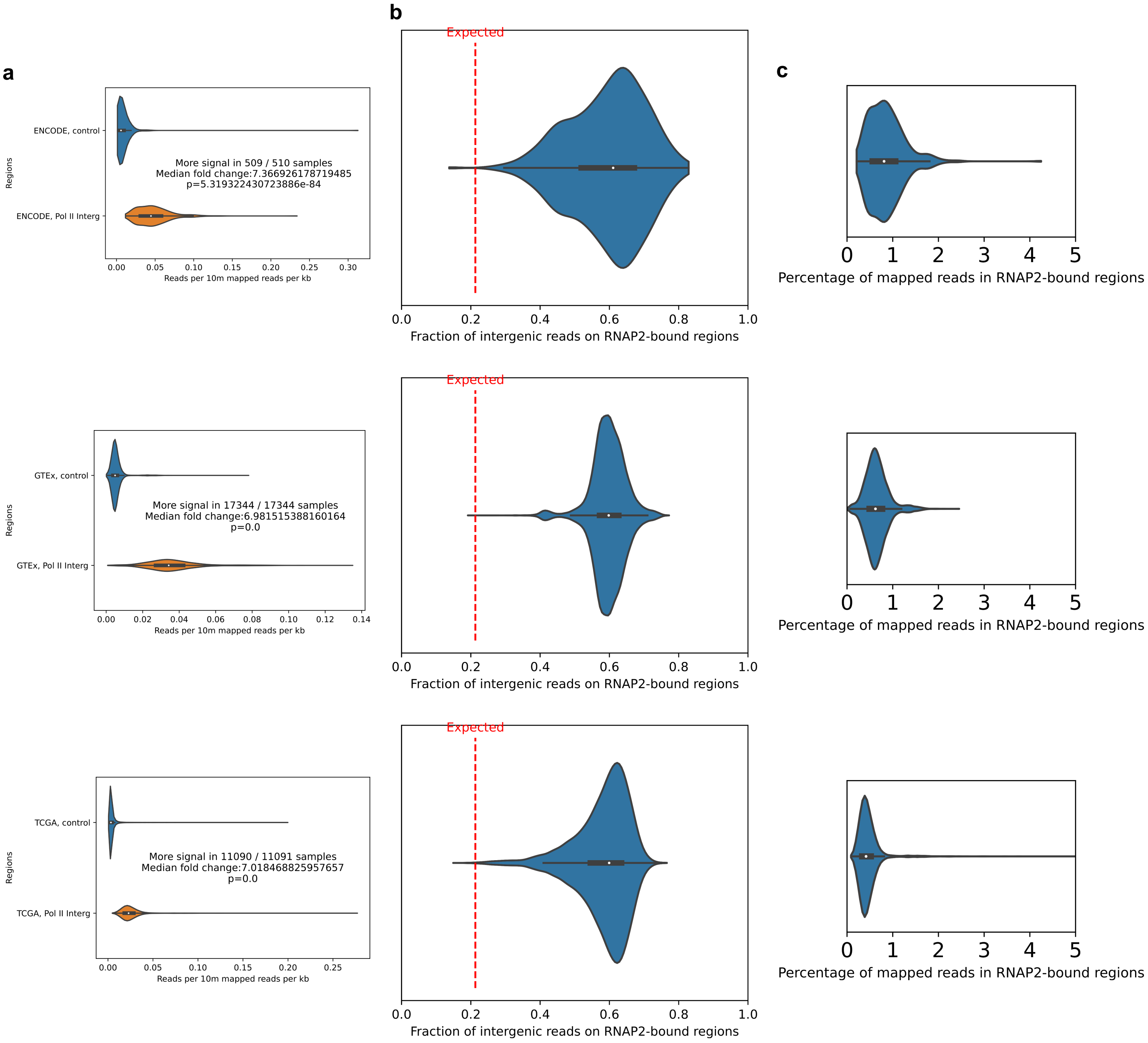


**Figure S8 : RNAP2-bound regions captures a majority of intergenic transcriptional signal**

**a.** Distribution of the average number of reads per RNAP2-bound region or control regions (non RNAP2-bound region, all 1kb binned intergenic regions), across samples (p-values from Mann-Whitney U-test). **b.** Distribution of the fraction of intergenic reads captured by RNAP2-bound regions across samples. Dashed line indicates expected value due to RNAP2-bound regions coverage. **c.** Distristribution of the percentage of total mapped reads captured by RNAP2-bound regions. From top to bottom : ENCODE, GTEx, TCGA rna-seq datasets.

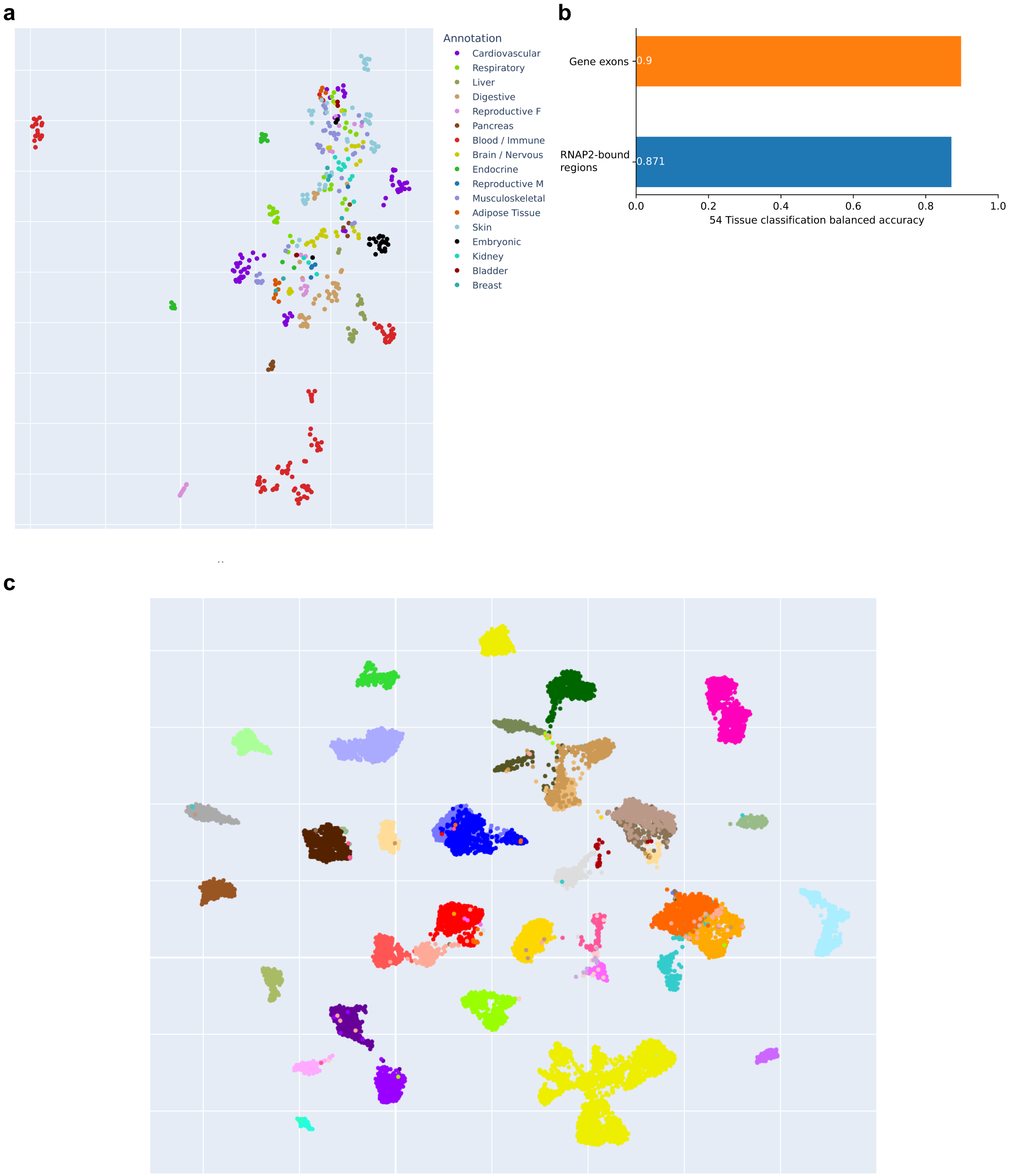


**Figure S9 : Intergenic transcription by itself is sufficient to characterise biological conditions**

**a.** UMAP of ENCODE total RNA-seq samples using rna-seq signal at RNAP2-bound regions. **b.** UMAP of GTEx RNA-seq samples using rna-seq signal at genes exons. **c.** KNN (5 NN, Pearson correlation as metric) classification balanced accuracy using either Gene expression or RNAP2 signal as input.

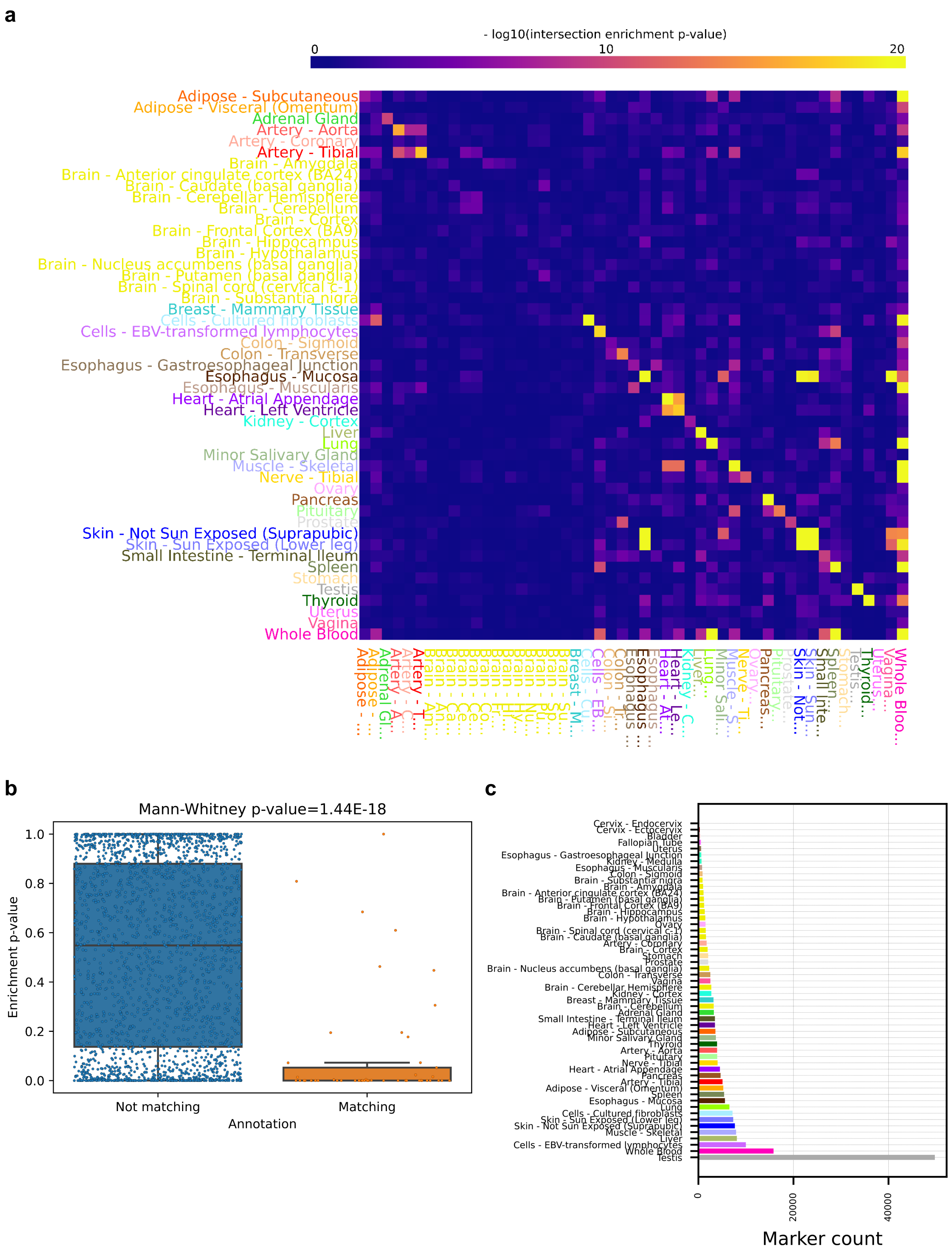


**Figure S10 : Tissue-specific regulatory variants are enriched within tissue-specific Intergenic transcripts**

**a.** Intersection enrichment heatmap between tissue-specific eQTLs (rows) and tissue-specific RNA-seq over-expressed marker RNAP2-bound regions. **b.** Distributions of tissue-matching (i.e. “Artery - Aorta” vs “Artery - Aorta”) and non-matching (i.e. “Artery - Aorta” vs “Artery -

Coronary”,“Liver”...) enrichment p-values between tissue-specific eQTLs (rows) and tissue-specific RNA-seq over-expressed markers. **c.** Number of overexpressed RNAP2-bound regions per tissue.

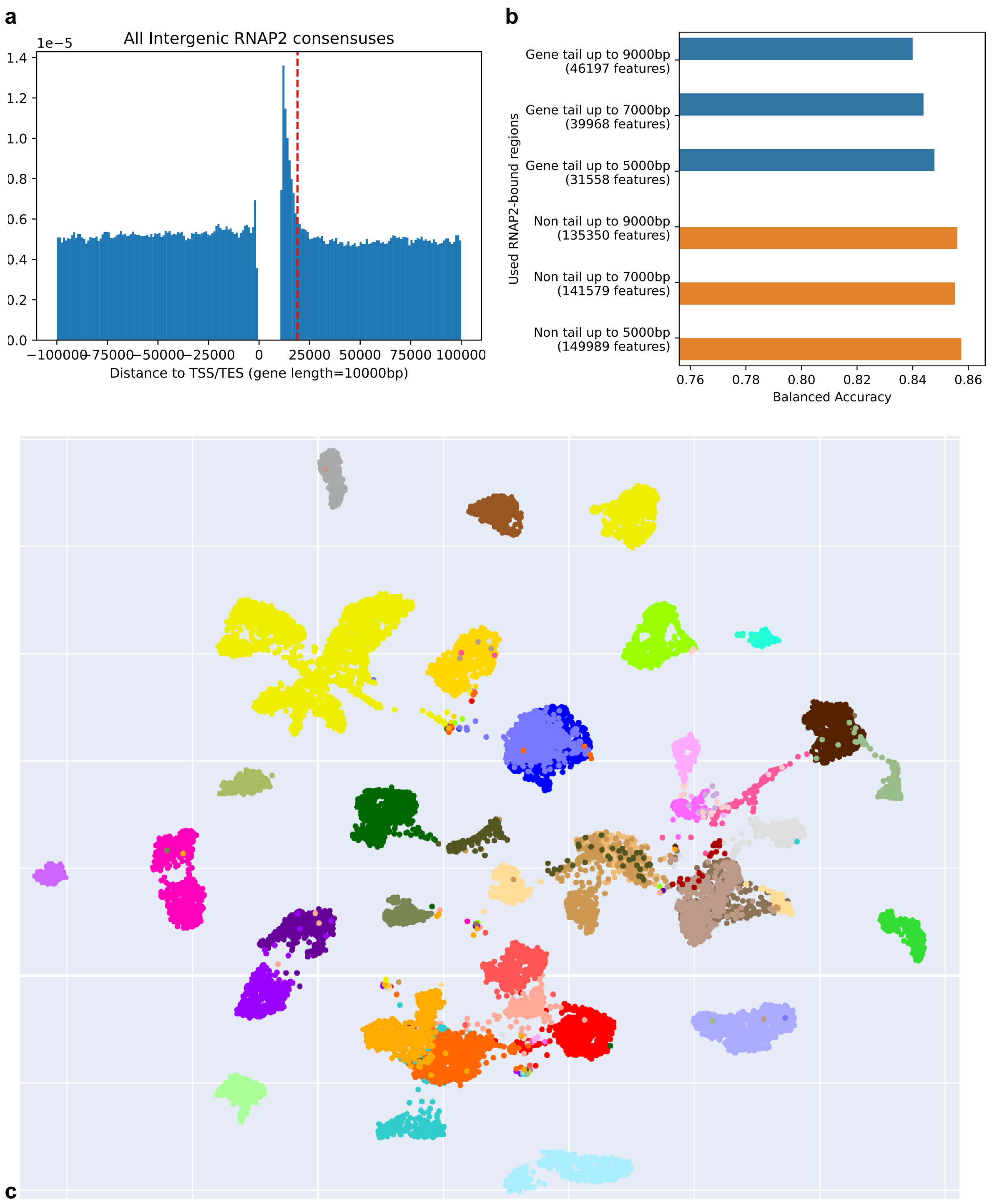


**Figure S11 : The intergenic transcriptional signal is not driven by end-of-gene transcription**

**a.** Distributions of RNAP2 consensus centroids relative to protein coding genes (gene length standardised to 10kb), red line indicates 9kb after Transcription End Site. **b.** KNN (5 NN, Pearson correlation as metric) classification balanced accuracy using different subsets of intergenic RNAP2-bound regions for classification. **c.** UMAP of GTEx RNA-seq samples using RNA-seq signal at intergenic RNAP2, excluding those located at less than 9,000bp of a Transcription End Site.

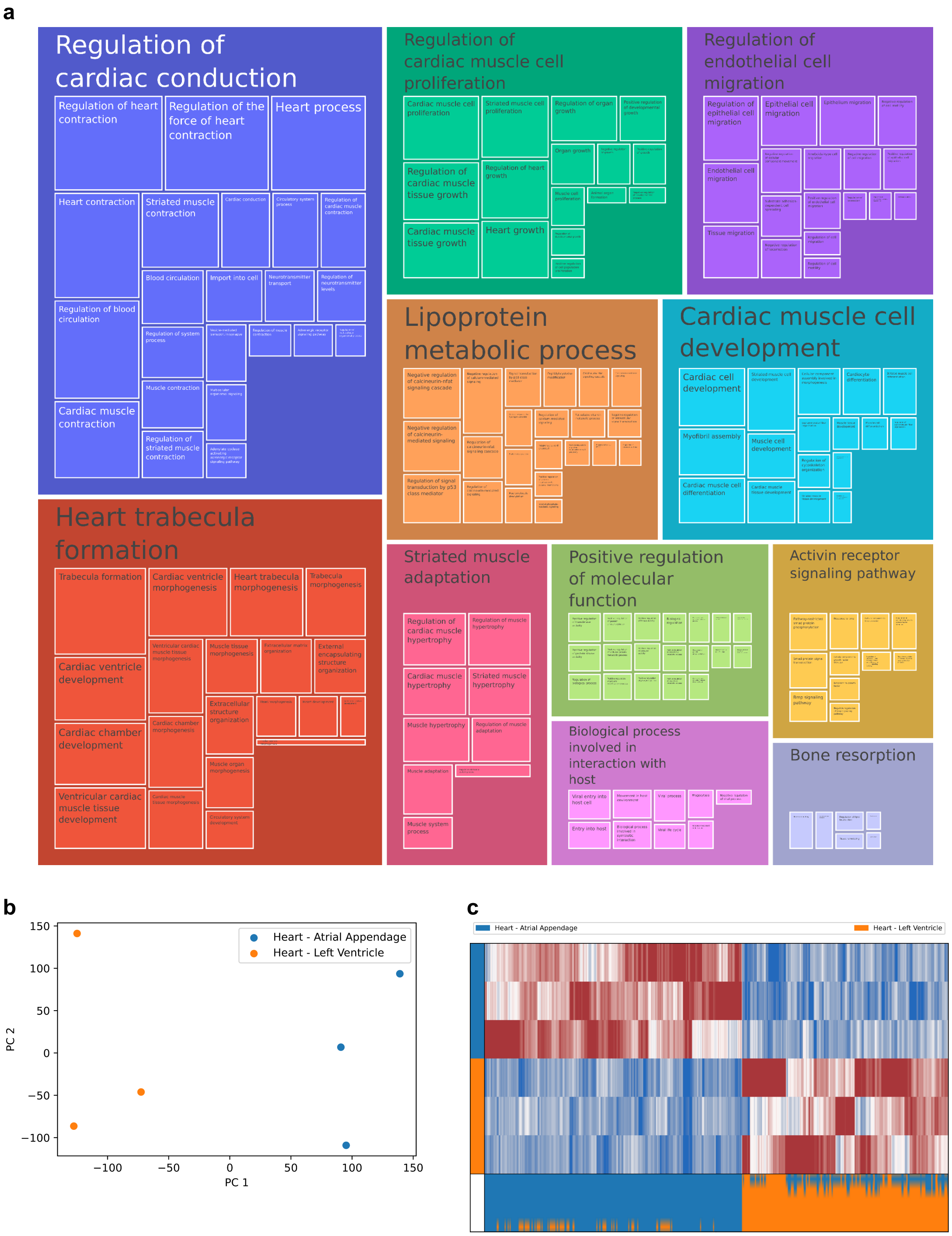


**Figure S12 : Differentially expressed RNAP2-bound regions can be detected at smaller sample sizes.**

**a.** Clustered GO terms enrichments for genes nearby RNAP2-bound regions differentially expressed between 'Heart - Atrial Appendage' and 'Heart - Left Ventricle' tissues from GTEx in a downsampled n=3 comparison (methods). **b.** First two principal components of RNAP2-bound region expression in a downsampled n=3 comparison. **c.** Heatmap of the Pearson residuals of DE RNAP2-bound regions in a n=3 comparison.

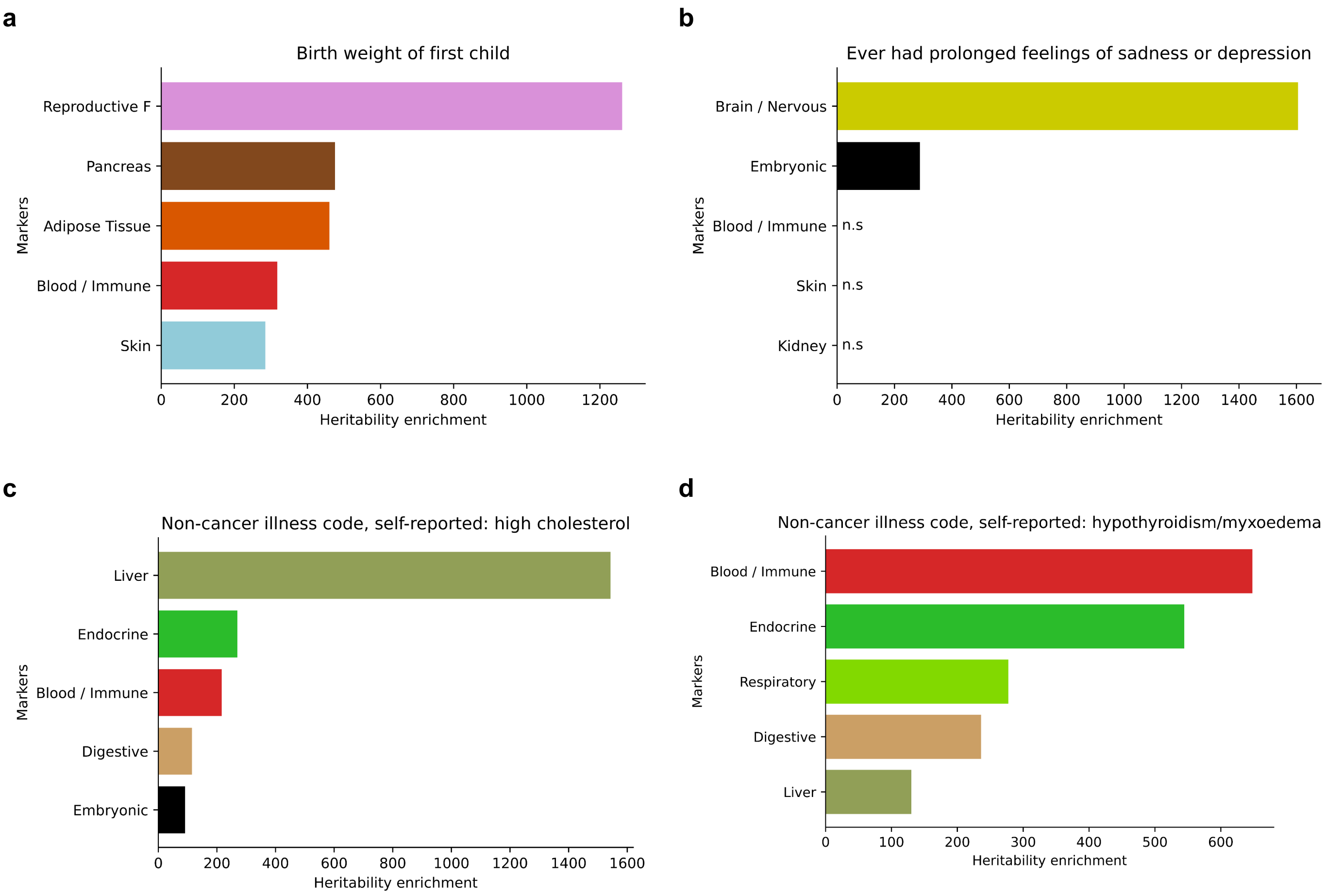


**Figure S13 : Per-biotype robustly over-expressed markers display meaningful disease-associated heritability enrichments**

Top 5 heritability enrichment over robust tissue-specific over-expressed markers (Methods), for 4 disease-associated UK Biobank GWAS traits. All enrichments are statistically significant unless otherwise mentioned.

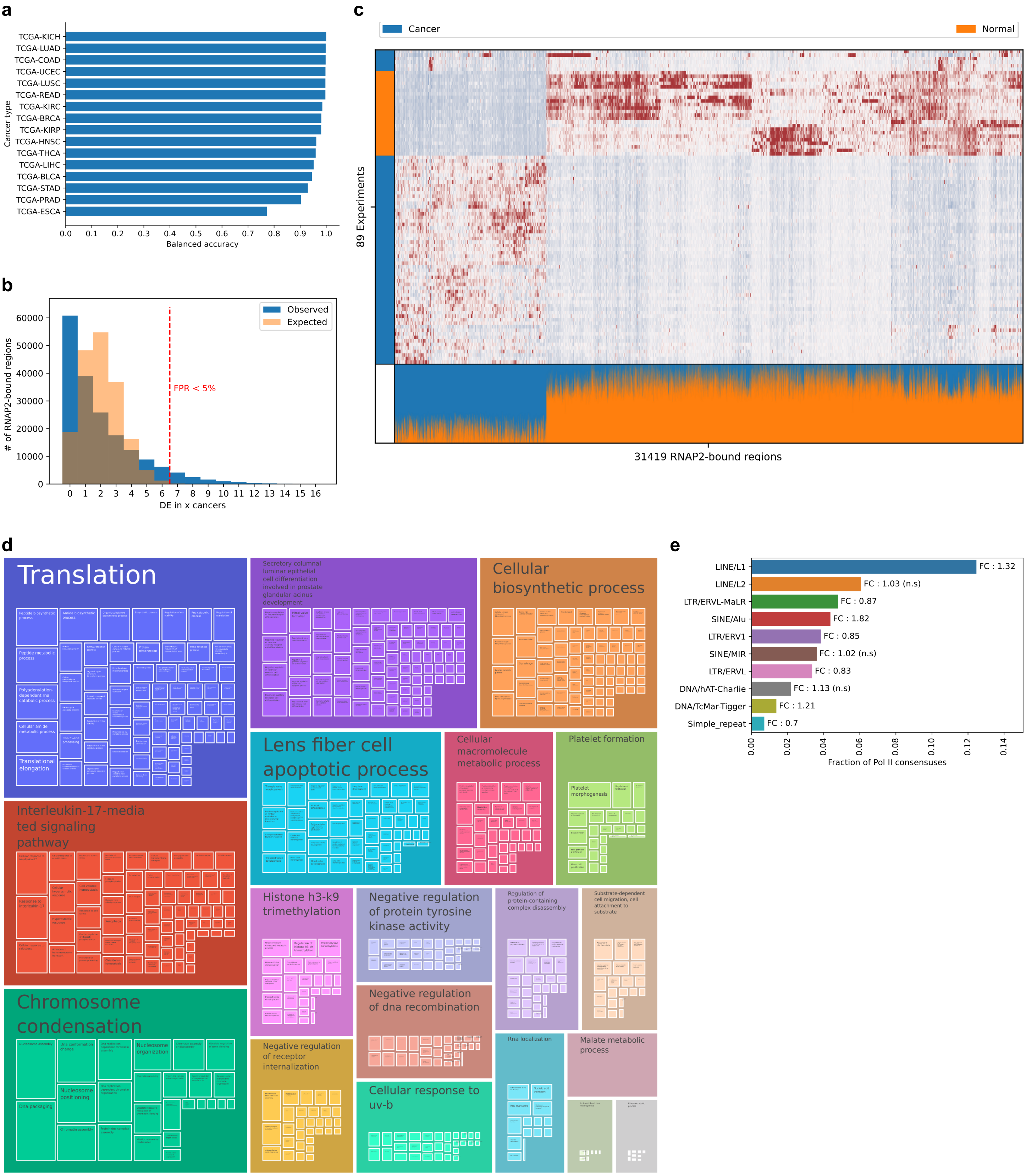
a
TCGA-KICH
TCGA-LUAD
TCGA-COAD
TCGA-UCEC
TCGA-LUSC
TCGA-READ
TCGA-KIRC
TCGA-BRCA
TCGA-KIRP
TCGA-HNSC
TCGA-THCA
TCGA-LIHC
TCGA-BLCA
TCGA-STAD
TCGA-PRAD
TCGA-ESCA
Cancer type
Balanced accuracy
b
Observed
Expected
FPR < 5%
### of RNAP2-bound regions
DE in x cancers
c
Cancer
Normal
89 Experiments
31419 RNAP2-bound regions
d
Translation
Secretory columnar luminar epithelial cell differentiation involved in prostate glandular acinus development
Cellular biosynthetic process
Interleukin-17-mediated signaling pathway
Lens fiber cell apoptotic process
Cellular macromolecule metabolic process
Platelet formation
Chromosome condensation
Histone h3-k9 trimethylation
Negative regulation of protein tyrosine kinase activity
Regulation of protein-containing complex disassembly
Substrate-dependent cell migration, cell attachment to substrate
Negative regulation of dna recombination
Negative regulation of receptor internalization
Cellular response to uv-b
Rna localization
Malate metabolic process
e
LINE/L1
FC : 1.32
LINE/L2
FC : 1.03 (n.s)
LTR/ERVL-MaLR
FC : 0.87
SINE/Alu
FC : 1.82
LTR/ERV1
FC : 0.85
SINE/MIR
FC : 1.02 (n.s)
LTR/ERVL
FC : 0.83
DNA/hAT-Charlie
FC : 1.13 (n.s)
DNA/TcMar-Tigger
FC : 1.21
Simple_repeat
FC : 0.7
Fraction of Pol II consensuses

**Figure S14 : Non coding transcription captured at RNAP2-bound regions discriminates normal and tumour tissues.**

**a.** Balanced accuracy of per cancer Gradient Boosted Decision Tree predictors of tissue state (Normal vs Tumour). **b.** Distributions of the number of cancers of RNAP2-bound region is DE in, as observed in the TCGA dataset, and by cancer-wise random permutations. Dashed line indicates the threshold at which less than 5% of observed DE RNAP2-bound regions are DE in more cancer than expected by chance. **c.** Heatmap of Pearson residuals (clipped at +-3) of DE RNAP2-bound regions in the Kidney Chromophobe Carcinoma dataset (KICH). Red = strongly expressed. Bottom part of the heatmap represents the fraction of normalised reads belonging to either class in each RNAP2-bound region (weighted by class imbalance). **d.** Clustered GO terms enrichments for genes nearby RNAP2-bound regions DE in 7 or more cancers (see methods). **e.** Fraction of DE RNAP2-bound regions in 7 or more cancers intersected for the top ten most intersected repeat families. FC corresponds to fold change enrichment versus random regions (see methods). All results are statistically significant unless otherwise mentioned.

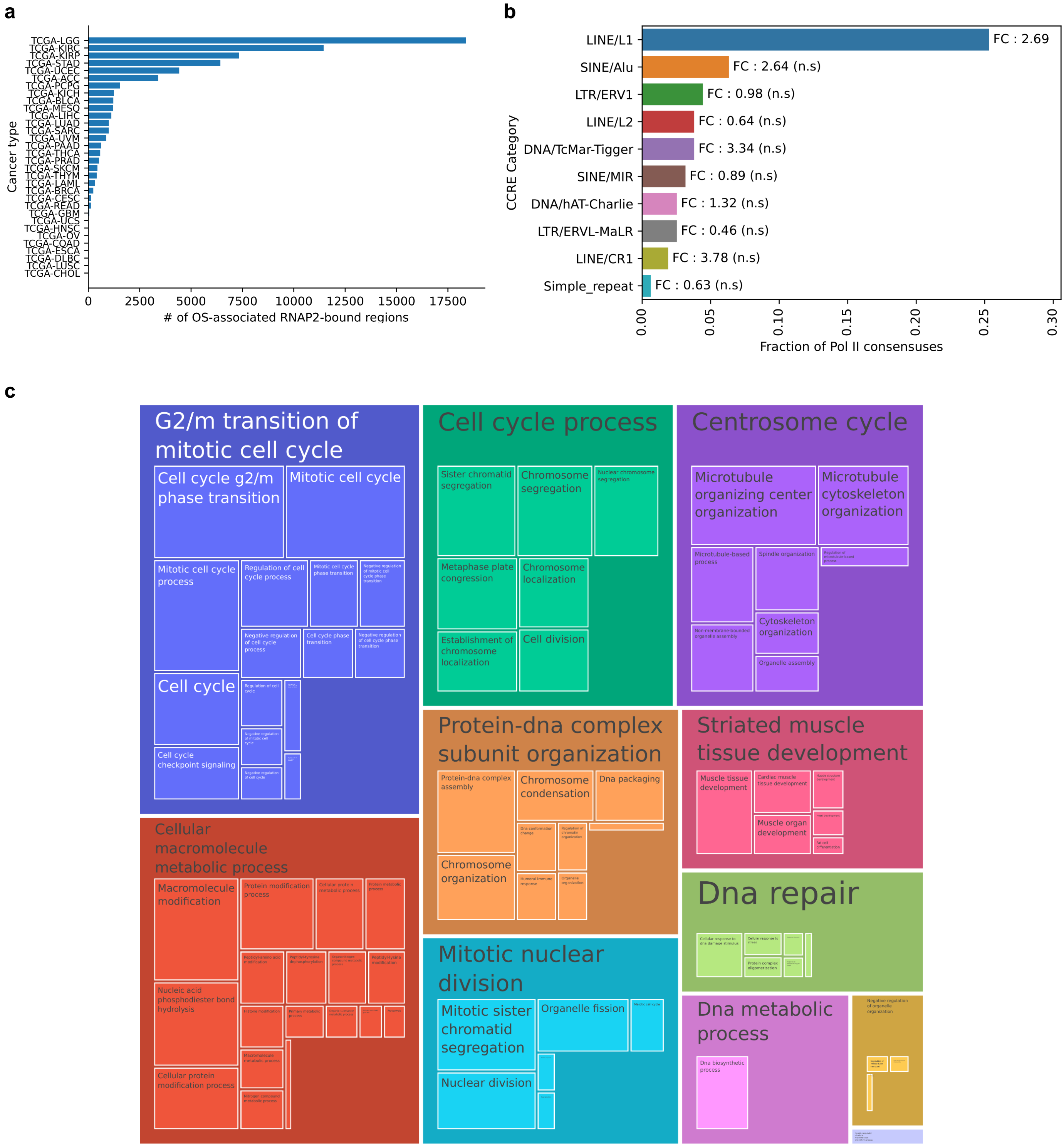


**Figure S15 : Non coding transcription captured at RNAP2-bound regions is prognostic of the patient's survival.**

**a.** Number of OS-associated RNAP2-bound regions for each TCGA cancer. **b.** Fraction of OS-associated RNAP2-bound regions in 5 or more cancers intersected for the top ten most intersected repeat families. FC corresponds to fold change enrichment versus random regions (see methods). All results are statistically significant unless otherwise mentioned. **c.** Clustered GO terms enrichments for genes nearby RNAP2-bound regions OS-associated in 5 or more cancers (see methods).
